# Confinement by liquid-liquid interface replicates *in vivo* neutrophil deformations and elicits bleb based migration

**DOI:** 10.1101/2023.06.14.544898

**Authors:** Jonathan H. Schrope, Adam Horn, Kaitlyn Lazorchak, Clyde W. Tinnen, Jack J Stevens, Mehtab Farooqui, Tanner Robertson, Jiayi Li, David Bennin, Terry Juang, Adeel Ahmed, Chao Li, Anna Huttenlocher, David J Beebe

## Abstract

Leukocytes navigate through interstitial spaces resulting in deformation of both the motile leukocytes and surrounding cells. Creating an *in vitro* system that models the deformable cellular environment encountered *in vivo* has been challenging. Here, we engineer microchannels with a liquid-liquid interface that exerts confining pressures (200-3000 Pa) similar to cells in tissues, and, thus, is deformable by cell generated forces. Consequently, the balance between migratory cell-generated and interfacial pressures determines the degree of confinement. Pioneer cells that first contact the interfacial barrier require greater deformation forces to forge a path for migration, and as a result migrate slower than trailing cells. Critically, resistive pressures are tunable by controlling the curvature of the liquid interface, which regulates motility. By granting cells autonomy in determining their confinement, and tuning environmental resistance, interfacial deformations are made to match those of surrounding cells *in vivo* during interstitial neutrophil migration in a larval zebrafish model. We discover that, in this context, neutrophils employ a bleb-based mechanism of force generation to deform a barrier exerting cell-scale confining pressures.

**Significance Statement:** Immune cells sense physical forces provided by surrounding cellular tissues to regulate their motility. Here, we introduce the use of liquid-liquid interfaces to model forces exerted by surrounding cells during interstitial motility *in vivo*. Neutrophils interacting with the interface employ a bleb-based mechanism of force generation to induce interfacial deformation. This work furthers our understanding of the mechanisms employed by immune cells to traverse through deformable barriers akin to cells in the body, and introduces a pioneering technology enabling the study of cell interaction with soft materials.

## Introduction

Precise regulation of leukocyte migration to sites of infection, malignancy, and injury is required for successful mounting and resolution of the immune response (1–4). Leukocyte motility is tightly regulated *in vivo* by a combination of chemical gradients and local mechanical cues arising from the surrounding physical environment (3–6). After exiting circulation, leukocytes migrate through tissue interstitial spaces and generate forces to overcome resistive forces applied by surrounding cells (3, 4, 6, 7). The balance between these leukocyte-generated and resisting forces regulates cell morphology and motility (4, 6). Existing *in vitro* migration platforms have uncovered that the degree of mechanical confinement and resulting variations in local cell shape regulate leukocyte speed, directionality, polarity, and migration mode (5, 8–12), thus implicating a role of local mechanical pressures in regulating the immune response. In the absence of confinement, leukocyte migration is largely mediated by actin polymerization-driven formation of a pseudopod at the leading edge (3, 4, 13, 14). However, under confinement, leukocytes can elect an alternative migration strategy driven by rear acto-myosin contractility driving membrane protrusions (blebs) through rapid fluctuations in intracellular hydrostatic pressure (12, 15–18).

It is well established that confining forces provided by the surrounding extracellular matrix (ECM) influence cell motility and effector function. However, our understanding of how interactions with surrounding cells themselves regulate motility is limited. The reasons for this are twofold; first, limitations in optical access of most *in vivo* models has thus far restricted capture of these single cell-cell physical interactions. Second, existing *in vitro* systems to study confined cell migration largely focus on cell-ECM interactions, or utilize materials orders of magnitude stiffer than cells themselves (5, 19–21) (**Fig. S1**). The development of ECM-mimics with tunable mechanical properties (22–24), and the ability to fabricate channels made of ECM-mimics (25–27), has yielded critical insights into how cell-matrix interactions modulate migration speed and directionality (28–31). However, such approaches neglect to model interactions with surrounding cells (stiffness ∼ 200-3000 Pa on single cell length scales (20) with very different mechanical properties than matrix fibers (∼300 MPa) (20). Alternative approaches utilize polymeric materials such as polyacrylamide, agarose, or polydimethylsiloxane (PDMS) based microchannels to study how controlled confinement of individual cells modulates cell traction (32, 33), polarity (34, 35), migration direction (10, 27, 36, 37), deformations (38), migration mode (18, 39), and speed (9, 40). However, the rigidity of these materials force cells into unnatural morphologies determined by pre-defined channel geometries. This has limited the ability to explore mechanisms of force generation employed by leukocytes to deform surrounding cells and forge paths through host tissues. Thus, there remains an unmet need for engineering an *in vitro* material interface that confines cells with physiologically relevant pressures to model physical interactions with neighboring cells and enable new insights into how interaction with soft materials regulates cell motility.

Here, we utilize differential surface patterning of a glass substrate to engineer microchannels on single cell length scales (i.e., channel height < 10 um) bound by a liquid-liquid interface. We examine interactions of primary neutrophils, the most abundant cells in the immune system and an exemplary model of amoeboid cell migration, with the liquid interface during chemotaxis. We find that the liquid-liquid interface exerts pressures comparable to cells themselves; sufficiently rigid for confinement, yet deformable in response to single cell generated forces. The result is greater autonomy granted to motile cells themselves to control the degree of confinement and thus morphology. Migrating neutrophils that reach the interface first (pioneer cells) require greater interfacial deformation than trailing cells and as a result migrate slower. Pioneer cells interacting with the interfacial barrier exhibit distinct morphological polarity characterized by frontward protrusions and rear-ward nuclear positioning. By careful tuning of the interfacial pressures resisting cell-generated forces, deformations of the interface are made to match those of epithelial cells during interstitial migration within a larval zebrafish model. While the majority of observations of bleb-based migration have occurred under rigid confinement, here we find that under *in vivo*-like confinement, neutrophils elect a bleb-based mechanism of force generation driven by rear contractility to deform the interface. Altogether, this work introduces a tunable material interface that replicates single cell-scale confining forces within a system accessible to a broad range of researchers to study how cells sense and respond to interactions with soft barriers to regulate motility.

## Results

### Exclusive Liquid Repellency (ELR) enables liquid microchannels of heights less than single cells

In this section, the surface engineering principles enabling construction of liquid channels of heights less than a single cell is described. Recently, groups have engineered liquid-walled microfluidic channels by either printing aqueous media onto a chemically homogeneous solid substrate under oil overlay (41–43), or by depositing media onto superhydrophilic regions of a chemically patterned (i.e., heterogeneous) substrate (44). In either case, interfacial forces within the three phase (solid-aqueous-oil) system resist aqueous fluid expansion and thus pin the media to user-controlled geometries. In such strategies, however, obtaining channel heights on single cell length scales (< 10 µm leukocyte diameter) is challenging, as the forces pinning the aqueous phase to the desired geometry do not reach the theoretical maximum possible within a three-phase liquid-liquid-solid system. Here, we leverage our recent discovery of exclusive liquid repellency (ELR) (45–47) to control interfacial geometry with the theoretical maximum of aqueous-confining forces, enabling engineering of conduit heights comparable to that of single cells (∼1-10 µm).

Engineering such channels is rooted in foundational principles of interfacial physics. The degree to which a solid surface repels a liquid within a three phase solid-liquid-liquid system is reflected by the Young’s contact angle (*θ*) that a liquid droplet (Liquid 1) makes on the surface (**Fig. 1A, top**). Young’s contact angle is determined by the balance between all interfacial energies (*γ*) present within the system. Meticulous engineering of this energy balance results in a surface exhibiting exclusive liquid repellency (ELR), that is, complete repulsion of a liquid, resulting in a contact angle of 180° (45, 46). Here, we differentially pattern a glass substrate to exhibit regions of ELR to aqueous media (Liquid 1) in the presence of silicon oil (Liquid 2) (i.e., Aqueous Repellent Surface (ARS)) (**Fig. 1A, middle**), or regions of ELR to silicon oil (Liquid 1) in the presence of media (Liquid 2) (i.e., Oil Repellent Surface (ORS)) (**Fig. 1A, bottom**). Controlling surface patterning of ARS and ORS regions to specific geometries (**Fig. S2**) results in a double-ELR system that provides the theoretical maximum of interfacial forces to pin both media and oil onto their preferred surface. Importantly, this enables the construction of channels bound by a liquid-liquid interface with heights less than single cell diameter (< 10 µm) (**Fig. 1B**). Media is deposited onto the pre-patterned surface by “sweeping” a hanging droplet of media across the substrate with a pipette to construct an array of > 50 channels on a standard microscope slide within seconds (**Fig. 1C**). Here, we implement a simple pattern that results in two hemispheric droplets connected by a single, linear, liquid channel (**Fig. 1D-E**). However the method allows for creation of an endless number of configurations (**Fig. S3**).

**Figure 1.**
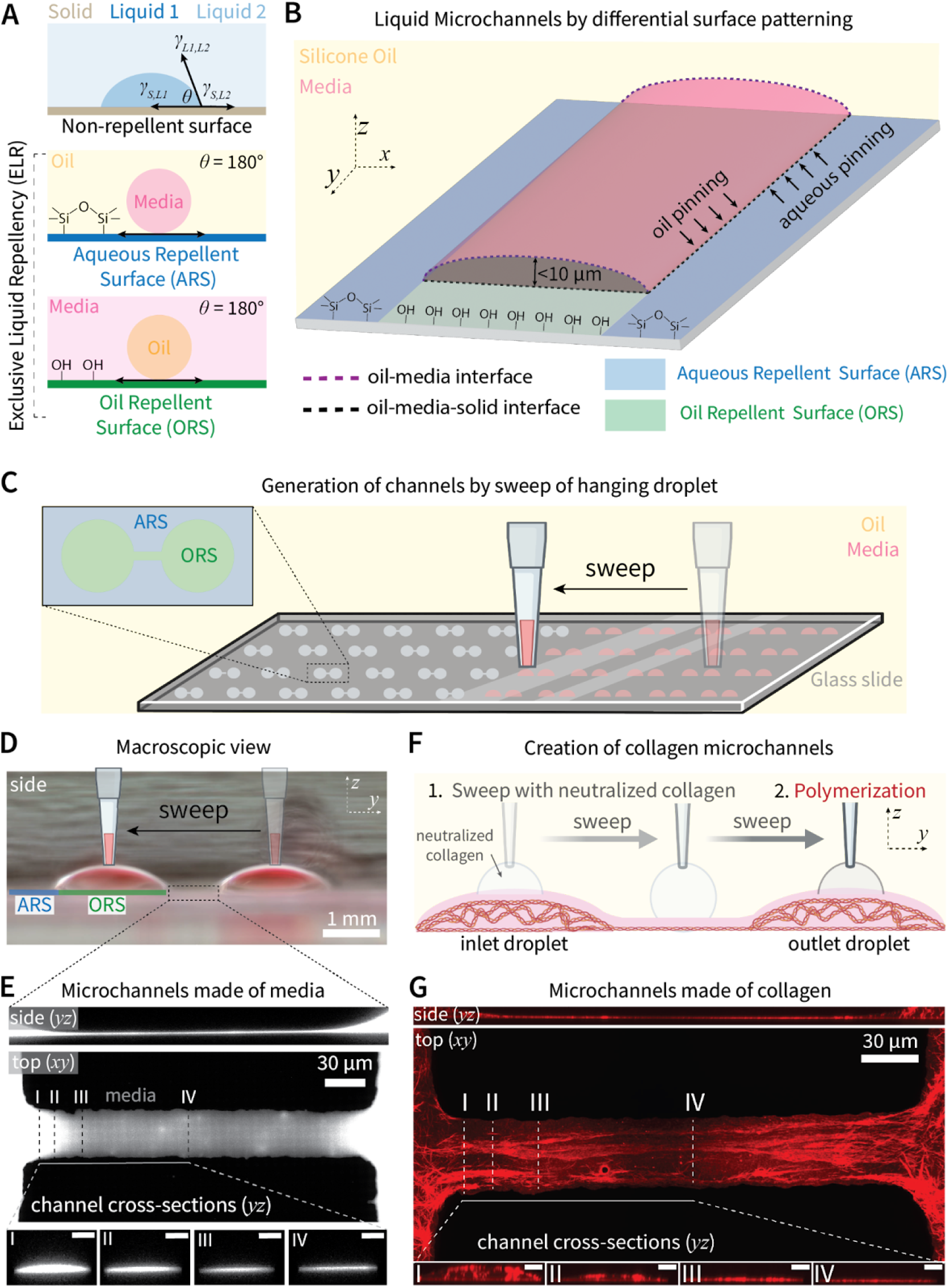
ELR-enabled construction of liquid microchannels on single cell length scales. **A)** Schematic depicting Young’s contact angle which describes the degree of liquid repulsion by a solid surface in a three phase (Solid-Liquid-Liquid) system. Controlling surface chemistry enables generation of surfaces exhibiting exclusive liquid repellency (ELR), characterized by complete liquid repulsion yielding a contact angle of 180°. Such surfaces can be Aqueous Repellent or Oil Repellent Surfaces (ARS, ORS respectively). **B)** Schematic depicting differential surface patterning of a glass substrate to exhibit ARS and ORS regions to create a double-ELR system. This enables creation of liquid channels with the maximum possible interfacial forces pinning both aqueous liquid and oil onto their preferred surface, enabling channel heights < 10 µm. **C)** Schematic of channel construction by “sweeping” a hanging droplet of aqueous media across ORS regions to form two hemispheric droplets and a connecting channel (> 50 on standard microscope slide). **D)** Macroscale side view of a media channel after sweep and addition of 1 µL of culture media to each droplet to improve visualization. **E)** Confocal image of a culture media channel with height < 1 µm (cross-sectional scale bars represent 3 µm). The channel height tapers at the entrance to a relatively consistent height (points III-IV) until a gradual, symmetric height increase towards the channel exit (right side). **F)** Schematic depicting incorporation of fibrous collagen into microchannels by performing the sweep with neutralized collagen then allowing polymerization. **G)** Representative confocal image of collagen-coated channel (cross-sectional scale bars represent 3 µm).

While collagen is the most abundant extracellular matrix (ECM) protein in the body and provides physical and chemical (48) cues to migrating cells, its high viscosity and rapid polymerization rate limits incorporation into traditional closed microchannels on sub-100 µm length scales. Existing methods often require external pumping of collagen into closed channels (49), decreasing throughput and increasing adoption barrier. Here, channels are constructed in open fluid, allowing for direct physical access so that performing the sweep technique with neutralized collagen yields microchannels on single cell length scales coated by fibrillar collagen (**Fig. 1F-G, Fig. S4**).

### The liquid-liquid interface confines, yet deforms in response to, immoble cells

In microchannel systems composed of rigid walls, the degree of confinement, and thus cell shape is determined entirely by the channel geometry. However during interstitial migration *in vivo*, leukocyte shape is determined by an equilibrium state between cell-generated and resistive pressures provided by the surrounding cellular environment. Thus, we sought to investigate whether the liquid-liquid interface 1) possesses sufficient rigidity to confine cells and 2) is “soft” enough to deform in response to single cell-scale forces (i.e. single cell stiffness). Primary human neutrophils were incorporated into the hanging droplet during the sweep and thus placed directly into channels of varying width (200 µm to 30 µm) (**Fig. 2A**). In the absence of cells, channel height is determined by an equilibrium state reflecting the pressure exerted by the aqueous phase itself; channel height (red line) increases with channel width (**Fig. 2C, Fig. S5A**). Cells placed into progressively smaller channels exhibited decreased cell height (blue line), implicating sufficient force applied by the interface to mechanically confine single cells (**Fig. 2B-C, Fig. S5B-F**). This demonstrates the ability to alter cell confinement by varying channel geometry. Notably, when channel height is significantly less than cell diameter (30 and 100 µm width channels), then cell height exceeds that of the channel alone (in the absence of cells). This implies that cells exert passive forces to deform the interface when initial channel height is sufficiently small. Indeed, confocal imaging along the length of a 30 µm width channel reveals that cells deform the interface from an initial equilibrium height (*h*_*0,equilib*_) to an increased equilibrium height (*h*_*cell,equilib*_) (**Fig. 2D**). Thus the interface is not a rigid barrier such as plastic or PDMS elastomer but is deformable by the presence of individual cells.

**Figure 2.**
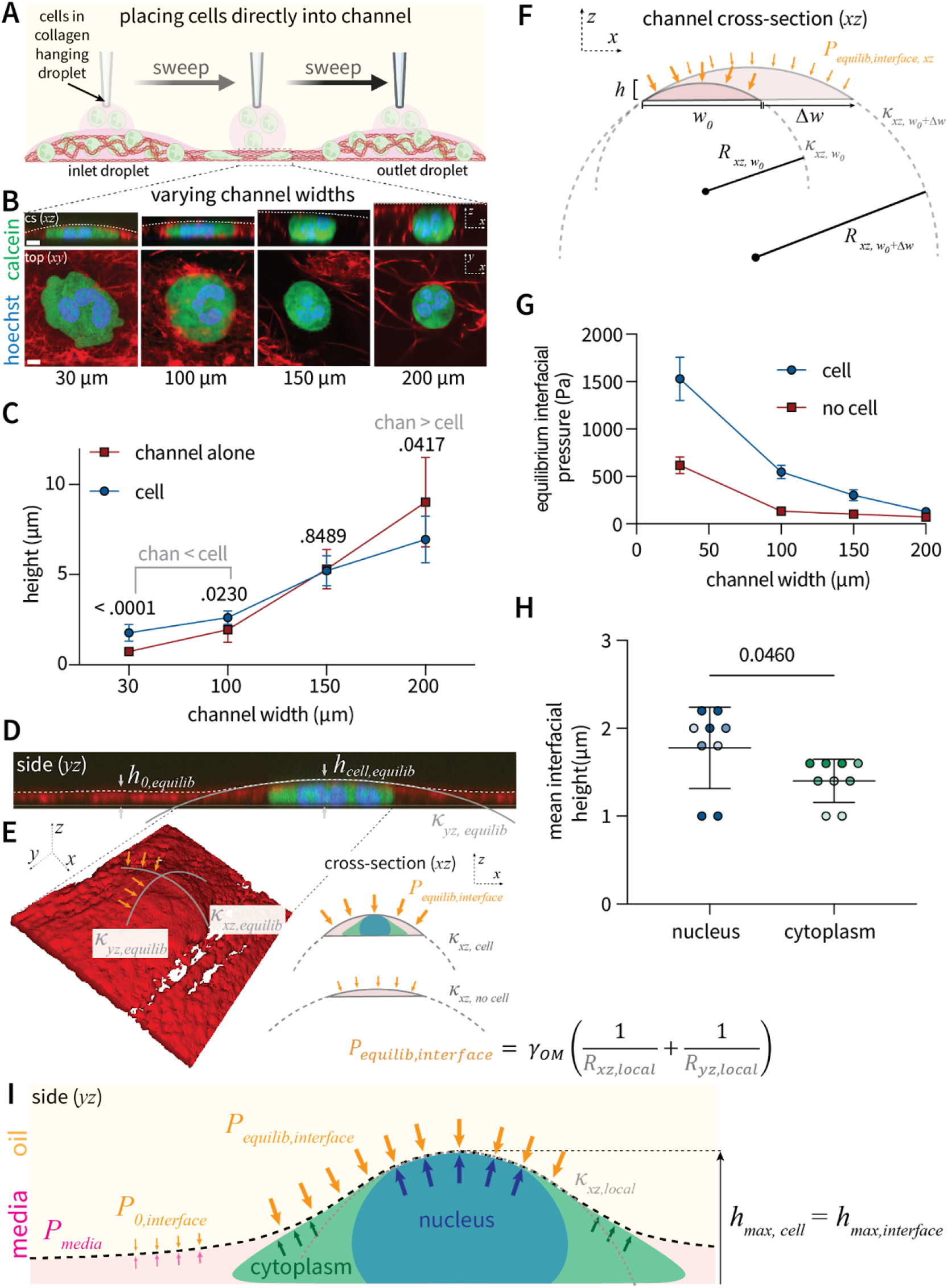
Tension of liquid-liquid interface exerts confining pressures on the scale of single cell stiffness. **A)** Schematic depicting direct addition of cells embedded in neutralized collagen into liquid microchannels by incorporation into the hanging droplet during the sweep process. **B)** Primary human neutrophils placed within channels of varying sizes (30-200 µm widths) exhibit differential degrees of cell confinement. **C)** Quantification of cell height as a function of channel width. Each point represents the mean cell (blue) or initial channel (red) height over three channel replicates for three different human donors. P-values represent differences between cell and channel height (blue vs red) determined by t-test. In channels of sufficiently low height (30 and 100 µm width), cell height exceeds that of channel height; on 200 µm width channels of height greater than cell diameter, cells exhibit a resting diameter less than that of channel height. **D)** Channel side view revealing that cells are confined by, yet deform the channel interface from *h*_*0,equilib*_ to *h*_*cell,equilib*_ to generate curvature (*κ*) in *yz* the plane. **E)** Confocal surface rendering depicting interfacial curvatures in the *xz* and *yz* planes resulting from cell-generated deformation. Central nuclear positioning in such immobile cells results in equivalent curvature in the *xz* and *yz* planes. **F)** Schematic defining the channel interfacial pressure (*P*_*equilib,interface*_), curvature (*κ*), and radius of curvature (*R*), in the *xz* plane. Increases in channel width alter interfacial pressure. **G)** Quantification of interfacial pressure (*P*_*equilib,interface*_) in the presence or absence of a cell as a function of channel width. Each point represents the mean of three independent channel replicates. **H)** The interfacial height is greatest over regions of the cell containing the nucleus compared to those containing only cytoplasm. Each point represents a cell measured on an independent channel replicate (n = 3), shade represents donor (n = 3). Statistical significance was determined by an unpaired, two-sample t-test assuming equal variance. **I)** Model schematic depicting equilibrium interfacial pressure defined by cell physical properties. In locations devoid of a cell, *P*_*equilib,interface*_ reflects the pressure exerted by the media itself. In the presence of a cell, *P*_*equilib,interface*_ is determined by the pressure exerted by the cell nucleus.

In contrast to rigid substrates with fixed stiffness dependent on material properties alone, the pressure exerted by a liquid-liquid interface is dependent both on interfacial components (i.e. oil and media) and the curvature of the interface. Thus we next sought to quantify the confining pressures exerted on cells by the interface. According to the Law of Laplace, the pressure exerted by any curved liquid-liquid interface is a function of the interfacial tension (material dependent constant; *γ*_*OM*_ ∼ 41.8 mN/m for water and silicone oil (46)) and the radii of interfacial curvature across all curved planes (side *yz* and cross-sectional *xz* in this case). Within this double ELR system that completely immobilizes the aqueous phase to the predefined channel width, deformations in channel height alter interfacial curvature (*κ*) and thus the reciprocal, radius of curvature (*R = 1/κ*) (**Fig. 2E**). The result is an equilibrium confining pressure that reflects the pressures exerted by cells themselves. Critically, interfacial curvature (and thus pressure) is controllable by altering channel width (**Fig. 2F**); interfacial pressures in the presence (blue) or absence (red) of cells decreases with increased channel width (**Fig. 2G**). This demonstrates the ability to control confining pressures by altering channel geometry. This mathematical framework would predict that the cell nucleus, the stiffest part of the cell, exerts the greatest pressure against the interface thus generating the greatest interfacial deformation. Indeed, the interfacial height is always greatest over the cell nucleus (**Fig. 2H**). Thus, in contrast to rigid-walled channels where cell confinement is purely a function of predefined geometrical constraints, these data support a model whereby the channels adopt a confining pressure reflecting an equilibrium defined by the physical properties of cells themselves (**Fig. 2I**).

### Neutrophil motility is dependent on required deformation of the interface

Given that the interface exerts confining pressures comparable to cells themselves, we sought to examine how interaction with the deformable barrier regulates cell motility. Creation of channels followed by sequential addition of neutrophils to the inlet droplet opposite from chemoattractant in the outlet droplet establishes a chemotaxis assay under confinement (**Fig. 3A, Fig S6A-E, Movie S1**). In this assay, cells occupy a layer of culture media between the collagen layer and oil-media interface, thus chemotaxis would require active deformation of the interface to allow space for migration (**Fig. 3B**). The initial cells to enter the channel, denoted as “pioneer” cells, migrate slowly, followed by “trailing” cells that migrate rapidly (**Fig. 3C-D**). Confocal imaging reveals differential interfacial heights before and after pioneer cell passage (**Fig. 3E-F**), establishing that pioneer cells actively deform the interface to allow space for migration. Therefore, while both populations are confined by the interface in the transverse (*z*) direction (**Fig. S6F**), pioneer cells must also interact with the interface in the axial (*y*) direction at the cell front. It is reasonable that this mechanical resistance is responsible for decreased pioneer cell speed. However, it is also possible that pioneer cell migration alters the chemical gradient in a manner to increase motility of trailing cells. To investigate this, we added a combination of fMLP and FITC dye of similar molecular weight (FITC 389.382 g/mol vs fMLP 437.56 g/mol) to visualize an approximate chemokine gradient during the course of migration (**Fig. S7A**). We found that pioneer cell passage did not significantly alter the chemical gradient (**Fig. S7B**). To further decouple positioning within the chemical gradient from required interaction with the interface, we analyzed a subset of trailing cells that migrate fast enough to catch up to and thus become pioneer cells. As these cells move through the stable global gradient, they significantly decrease in speed once they reach the interface (**Fig. 3G**). Taken together, these data indicate a purely mechanical mechanism whereby the interaction with (and deformation of) the interface restricts cell motility. Pioneer cells actively deform the channel interface, leaving decreased mechanical resistance for trailing cells which in turn migrate faster. From a technical standpoint, these results demonstrate the ability to decouple chemical cues such as position within a stable gradient, from mechanical cues of interaction with the interfacial barrier to isolate a role of physical cues on cell motility. Given the ability to tune pressures exerted by the interface, we next studied how alterations in the physical properties of the surrounding environment regulate cell motility.

**Figure 3.**
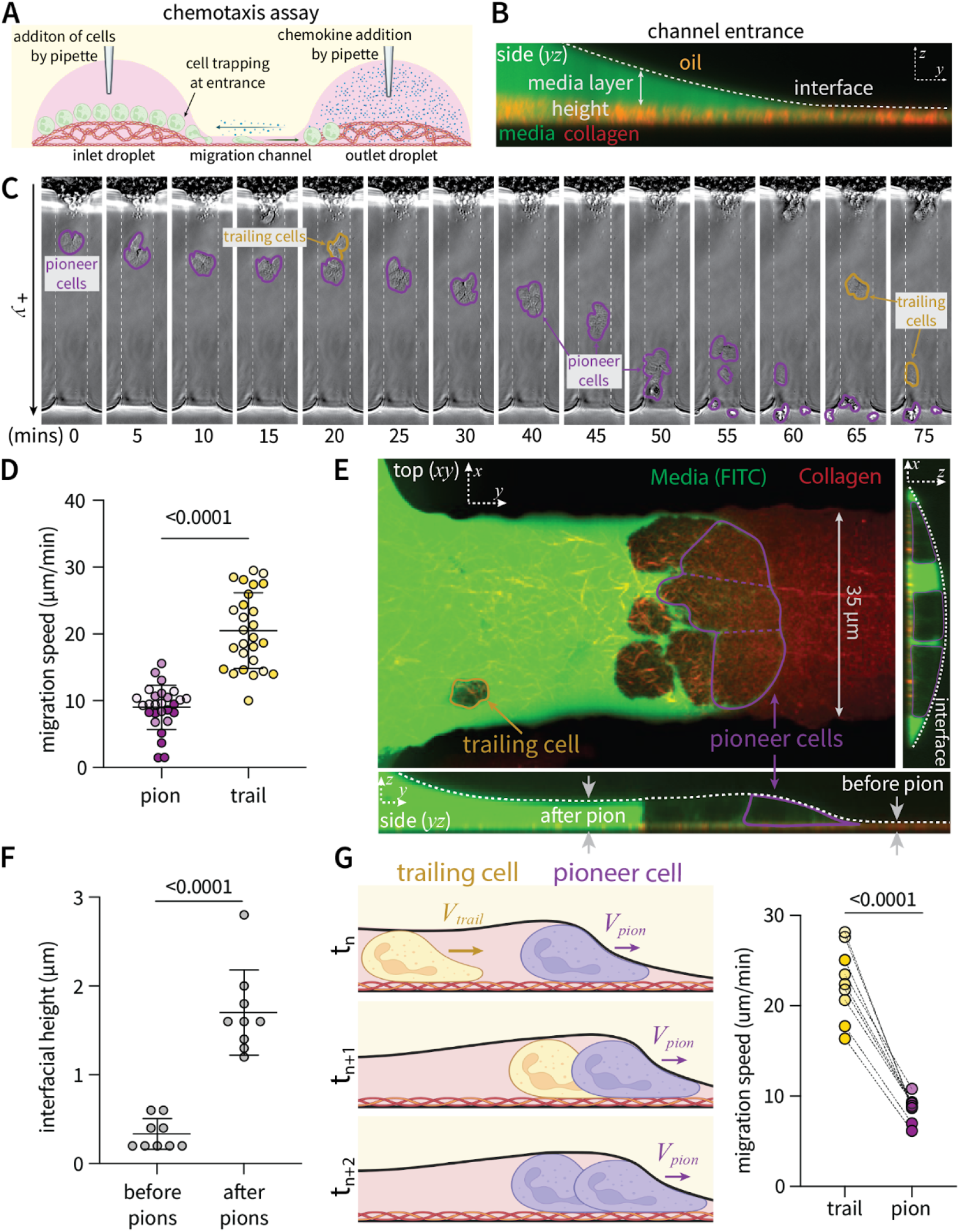
Migration speed is dependent on required deformation of interface. **A)** Schematic of migration experiment whereby collagen channels are created by the sweep technique followed by sequential addition of cells in the inlet droplet across from chemoattractant in the outlet droplet. **B)** Representative (confocal) side view of channel entrance. Cells are confined to the media layer between the collagen coating and oil-media interface. **C)** Timelapse images and **D)** corresponding quantification of migration speed for pioneer and trailing cells (n = 5 cells depicted by circles over 3 channel replicates and 3 independent human donors denoted by shade). **E)** Confocal image of pioneer cells deforming the interface (dashed white line). Media was visualized by addition of FITC dye. **F)** Quantification of interfacial heights before and after passage of pioneer cells. Each point represents an independent channel replicate (n = 10). **G)** Model schematic and corresponding quantification of migration speed for neutrophils that start as trailing cells but catch up to and become pioneer cells over time. Each data point represents the mean speed of a cell before and after the transition to becoming a pioneer (n = 3 cells observed on independent channel replicates over n = 3 independent human donors, represented by shade). Statistical significance was determined by either an unpaired (D,E) or paired (G) two-sample t-test assuming equal variance. All data gathered on 30 µm width channels.

### Liquid interface generates a tunable gradient of confinement that regulates motility

In contrast to immobile cells, migratory cells typically exhibit morphological polarity, largely dependent on nuclear positioning as they extend cytoplasmic protrusions at the leading edge to deform the surrounding environment. Rigid microchannels impose fixed environmental stiffness that is not responsive to cell-generated forces, so that all cell components (nucleus and cytoplasm) typically assume the entirety of available space within the channel. Here, we find that pioneer neutrophils exhibit rear-ward nuclear positioning and front-ward cytoplasmic protrusions (**Fig. 4A, Fig. S8A, Movie S2**). These regions deform the interface to different degrees; the nucleus provokes the maximal interfacial deformation, characterized by increased interfacial curvature (**Fig. 4B**) and thus maximal confining pressures, compared to frontward protrusions (**Fig. 4C**). Thus in contrast to immobile cells directly placed into channels, which exhibit symmetric deformations in the *xz* and *yz* planes, pioneer cells interacting with the interface generate spatially-variant radii of curvature across both planes. The result is a gradient of confining pressures across the length of the cell. This gradient is tunable by altering channel width (**Fig. S8B**). Increased width generates a more shallow gradient across cell length (*yz* plane), with less significant differences in confining pressure between the forward protrusion and nucleus-containing rear (**Fig. 4D**). Therefore tuning interface curvature (*xz* plane) controls cell shape (*yz* plane) during pioneer cell migration. Bulk analysis confirms that increased channel width results in decreased confining pressures over both the cell nucleus (**Fig. S8D**) and frontward protrusions (**Fig. S8E**). Thus akin to interstitial migration *in vivo*, the greater the force generated by the cell, the higher resistive pressure exerted by the surrounding elastic environment.

**Figure 4.**
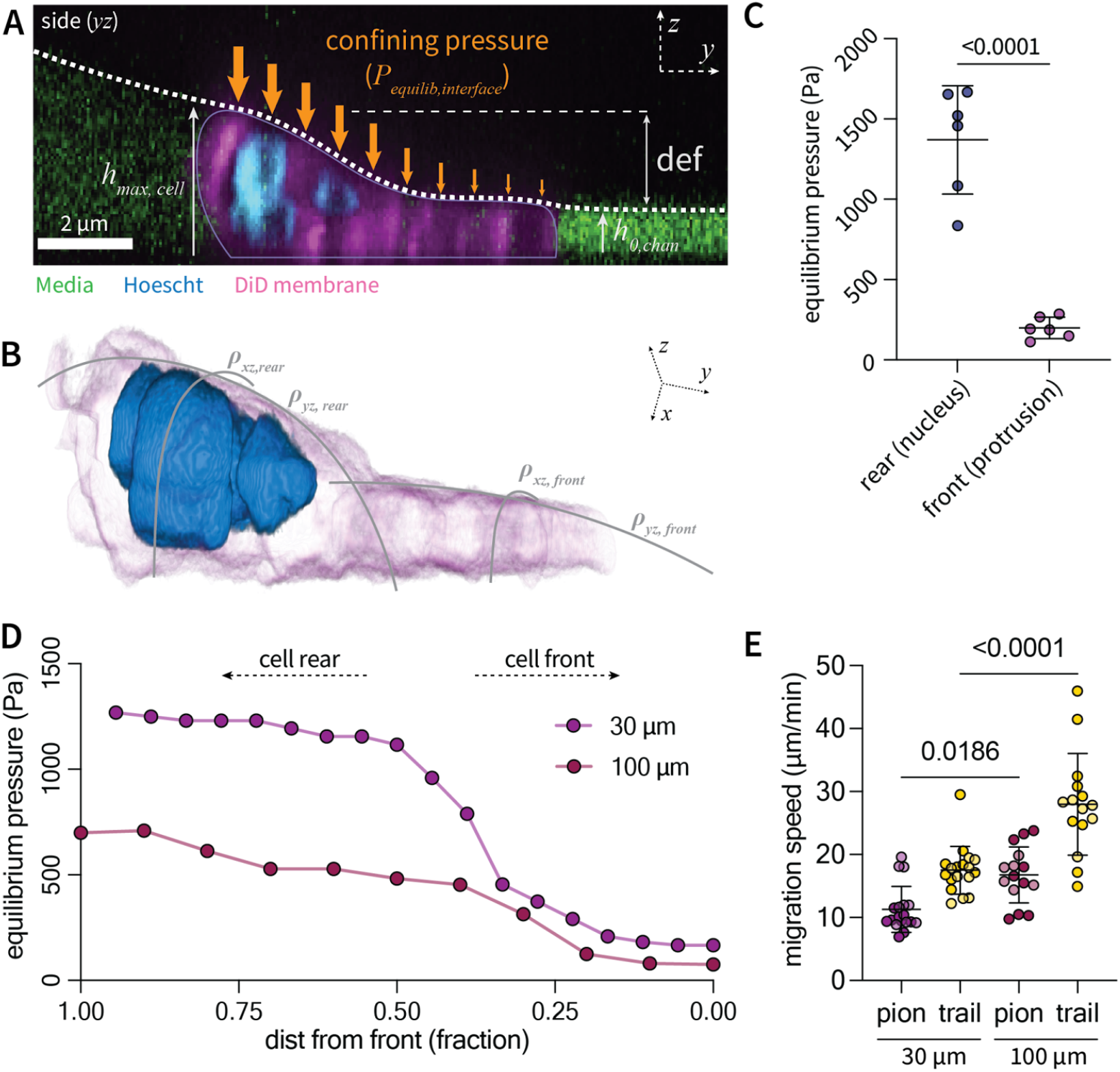
Liquid-liquid interface exerts gradient of confining pressure on migratory cells exhibiting morphological polarity. **A)** Representative side view of neutrophil migration within a 100 µm width channel exhibiting rear-ward nuclear positioning and a front-ward protrusion. This results in differential interfacial deformation and thus equilibrium confining pressure across the length of the cell (*yz* plane). **B)** Three-dimensional confocal reconstruction depicting differential interfacial curvature across cell length. **C)** Quantification of equilibrium confining pressure over the cell rear and front. Statistical significance determined by an unpaired, two-sample t-test assuming equal variance. **D)** Representative confining pressure gradients over the length of pioneer cells within channels of differential width (30 µm vs 100 µm). **E)** Cell speed as a function of channel width. Each point represents a single cell, data pooled over three independent channels and two independent donors (denoted by shade). Statistical significance determined by a one-way ANOVA with multiple comparisons between every group.

To explore how altering confining pressure gradients regulate motility, we engineered channels of varying width (30 vs 100 µm) and length (300 vs 500 µm, respectively) that exhibit different confining pressures yet similar chemical gradients (**Fig. S8B-C**). While the chemical gradient within 30 µm width channels was slightly steeper than that in 100 µm channels (typically increasing motility), cells migrated significantly slower, suggesting that increased confining pressures negatively impacts motility (**Fig. 4E**). Ultimately, these data indicate that the liquid-liquid interface is sensitive to differential pressures generated across a single migratory cell, and that cell morphology and motility is tunable by controlling channel geometry. These data would predict rearward nuclear positioning eliciting maximal surrounding cell deformation during passage between surrounding cells *in vivo*, which we demonstrate in a larval zebrafish model.

### Tuning interface pressures recapitulates surrounding cell deformations during interstitial migration *in vivo*

Migratory leukocytes sense membrane deformations to regulate motility and effector function (50–53). Membrane shape *in vivo* is determined by a balance between outward cell generated forces and resistive pressures arising from surrounding cells. Thus, the magnitude of surrounding cell deformations is a reflection of the relative force relationship between migratory and surrounding cells. We next explored whether tuning the resistive pressures applied by the interface could recapitulate relevant leukocyte morphology and surrounding cell deformations during interstitial migration *in vivo*.

Limitations in optical access of most *in vivo* models has hindered the ability to capture leukocyte physical interactions with surrounding cells during migration. Here, we leverage the genetic tractability and optical transparency of the larval zebrafish to capture deformations of neutrophils and surrounding keratinocytes during interstitial migration within the zebrafish epidermis (**Fig. 5A**). We performed high resolution imaging of spontaneous neutrophil motility within the epidermis of a dual transgenic larval zebrafish model expressing BFP in the neutrophil cytoplasm, mCherry in the nucleus and GFP in surrounding basal keratinocytes (*Tg(LyzC:TagBFP/LyzC:H2B-mCherry) x Tg(Krtt1c19e:acGFP)*). Such imaging reveals that neutrophils interact with and deform surrounding basal keratinocytes during interstitial migration (**Movie S3**). Neutrophils first extend a cytoplasmic protrusion to exert outward forces on surrounding keratinocytes, followed by a period of maximal keratinocyte deformation during passage of the cell body (**Fig. 5B**). As predicted within liquid channels, the nucleus is rear-ward oriented, and its passage induces maximal deformations of surrounding cells as the case of the liquid interface (**Fig. 5C**). Tuning the interfacial shape by controlling channel width (30 µm vs 100 µm) reproduces neutrophil protrusion width (**Fig. 5D**) and surrounding cell deformations during passage of the cell body (**Fig. 5E**) observed *in vivo*. These data indicate that tuning of baseline channel pressures reproduces relevant transient force balances during leukocyte interstitial migration *in vivo*, validating the relevance of this system to model physical cell-cell interactions during migration.

**Figure 5.**
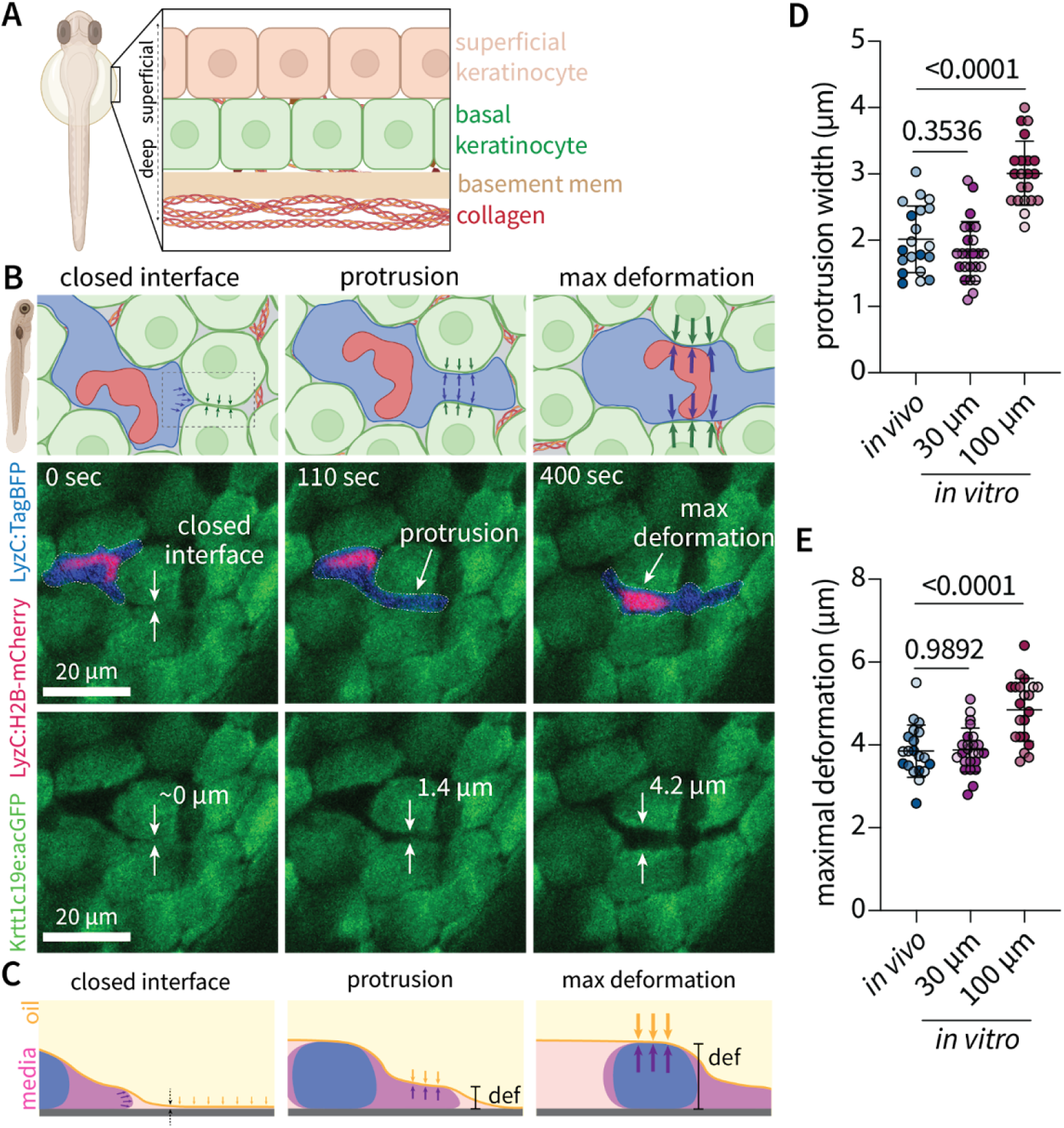
Tuning interfacial geometry recapitulates surrounding cell deformations during neutrophil migration *in vivo*. **A)** Schematic of skin layers in larval zebrafish three days post-fertilization. At this developmental stage, the epidermis consists of a single layer of both superficial (tan) and basal (green) keratinocytes. **B)** Schematic and corresponding time-lapse images depicting distinct phases of interstitial migration *in vivo* whereby neutrophils extend protrusions to deform surrounding basal keratinocytes (green) followed by passage of the cell body resulting in maximal deformation of surrounding cells. **C)** Schematic depicting interfacial deformation during protrusion extension and passage of the cell body *in vitro*. **D-E)** Tuning interfacial geometries recapitulates neutrophil protrusion width (D) and maximal surrounding cell deformations induced by the neutrophil cell body (E). Circles represent single cell passage events, shade represents independent biological replicates (n = 3), either a different fish clutch or human donor. Statistical significance was determined by a one-way ANOVA assuming equal variances with multiple comparisons to the *in vivo* group.

### Neutrophils employ a bleb-based mechanism of force generation to deform an interface exerting cell-scale confining pressures

We next sought to explore the mechanism of force generation that cells employ at the leading edge to deform the interface and generate space for migration. On two-dimensional substrates, leukocytes largely rely on polymerization of branched actin networks stabilized by actin-related protein 2/3 (Arp2/3) to extend protrusions at the leading edge. Recent studies have demonstrated that under rigid confinement, neutrophils can employ a bleb-based motility mechanism independent of actin polymerization at the leading edge (18), however it is not known how cells generate forces to deform a surrounding elastic environment exerting cell scale confining forces. Blebbing is driven by Rho-kinase dependent contraction at the rear driving increased intracellular hydraulic pressure. This drives detachment and rapid projection of the cell membrane through the F-actin cortex at the cell front (**Fig. 6A**). Blebs form on significantly faster timescales (< 2 µm/sec and < 10 sec lifespans) than lamellipodia (12). We investigated the hypothesis that interaction with the interface provoked a bleb-based mechanism of migration to generate outward deforming forces.

**Figure 6.**
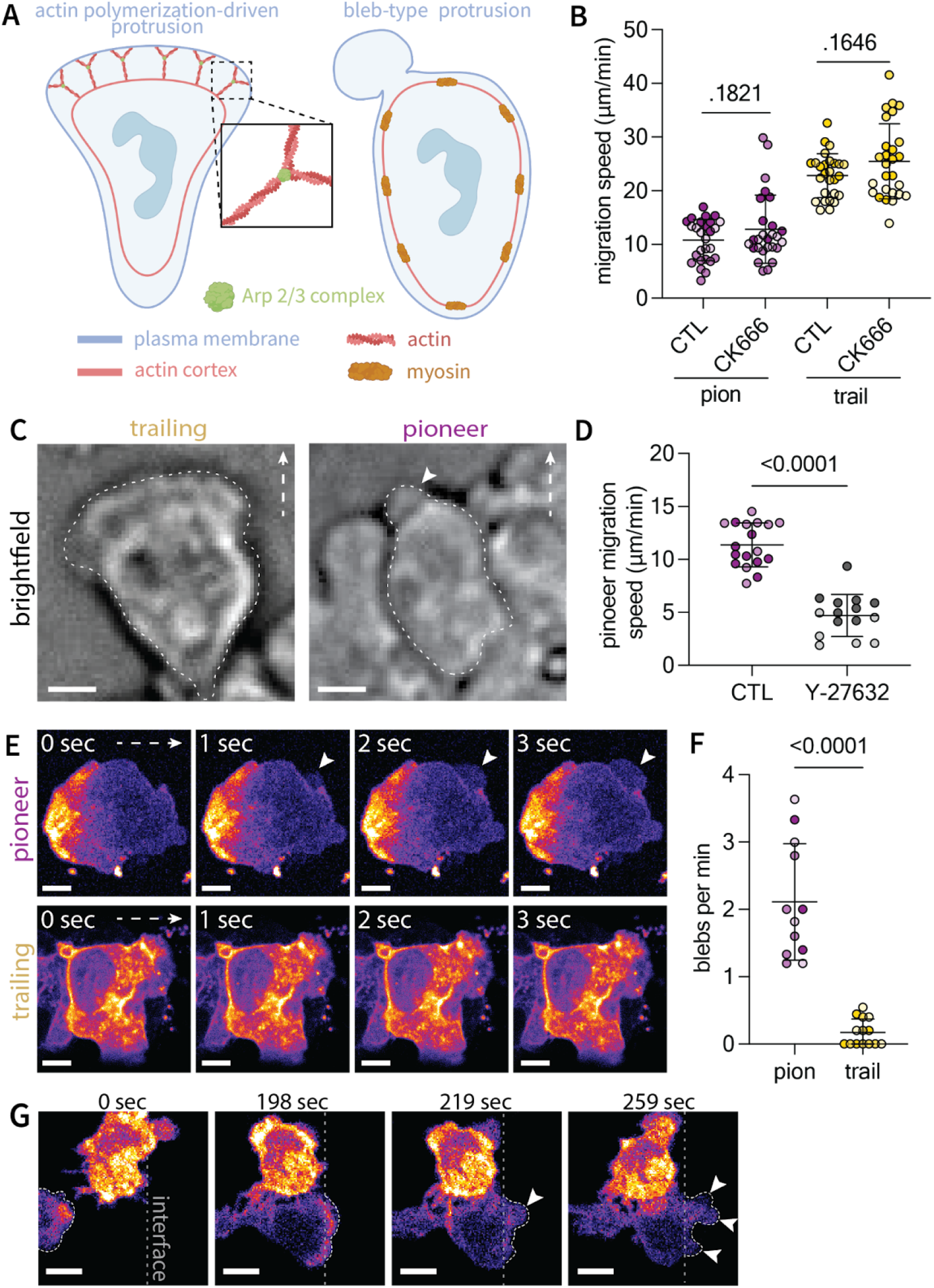
Neutrophils employ a bleb-based mechanism of force generation to deform the confining interface. **A)** Schematic of leukocyte migration modes, including actin polymerization-dependent pseudopod formation and bleb formation through rear contractility driving membrane extension. **B)** Treatment with Arp2/3 inhibitor CK666 does not impact migration speed within liquid channels. Circles represent mean track speed of individual cells pooled over three technical channel replicates; shade represents individual donor replicates (n = 3). **C)** Image of bleb-like protrusions formed by primary neutrophils compared to pseudopodia of trailing cells; scale bar represents 10 µm. **D)** Inhibition of rear contractility with Rho Kinase inhibitor Y-27632 decreases pioneer migration speed in primary neutrophils. White arrow denotes migration direction. Circles represent cell speeds pooled over three channel replicates; shade represents human donor replicate. **E)** Timelapse images of neutrophil-like PLB-985 cells expressing LifeAct-mRuby depicting bleb formation within pioneer cells and pseudopodia characteristic of trailing cells. White arrow denotes migration direction. Scale bar represents 5 µm. **F)** Quantification of blebs per minute for pioneer and trailing neutrophil-like PLB-985 cells. Each point represents a cell, data pooled over three independent biological replicates represented by shade. **G)** Trailing cells that reach the interface transition from pseudopodia to bleb-based protrusions (denoted by white arrows); scale bar represents 10 µm.

We first examined migration of primary neutrophils under inhibition of the Arp-2/3 complex using the small molecule CK666. While migration was inhibited within an established two-dimensional migration system (**Fig. S9A-B)**, it was unaffected under confinement within liquid channels (**Fig. 6B**). This suggests a migration mode independent of Arp-2/3 mediated actin polymerization at the leading edge. Intriguingly, rapid imaging (1 sec intervals) revealed sheet-like pseudopodial protrusions in trailing cells and formation of rapid, rounded protrusions that mimic the shape and kinetics of blebs in pioneer cells (**Fig 6C, Fig S9C, Movie S4**). These bleb-like structures are disproportionately formed by pioneer cells interacting with the interface compared to trailing cells (**Fig S9D**). Inhibition of rear contractility under treatment with Rho-Kinase inhibitor Y-27632 hinders the ability of pioneer cells to migrate through the interfacial barrier (**Fig. 6D**). Under such treatment, pioneer cell speed is sufficiently slow such that trailing cell motility is limited by that of pioneers and thus were excluded from analysis. Attempts at confirming blebbing in primary neutrophils by visualization of F-actin was limited by photobleaching of fluorescent actin probes and resistance to transfection. Thus, we imaged migration of neutrophil-like PLB-985 cells expressing LifeAct-mRuby. Pioneer neutrophil-like cells exhibited prominent blebbing at the leading edge compared to trailing cells that favored pseudopodial-based protrusions (**Fig. 6E-F, Fig. S9E, Movie S5-6**). Furthermore, trailing cells that eventually reach the interface to become pioneers transition from utilizing pseudopodia to bleb-based protrusions (**Fig. 6G, Movie S7**). Taken together, these data indicate a mechanism of bleb-based force generation at the leading edge to deform an interface exerting cell-scale confining pressures.

## Discussion

It is well established that cells sense physical properties of the surrounding tissue environment to regulate motility, oftentimes by sensing changes in cell membrane shape. This realization has predominantly relied on development of synthetic biomaterials in attempts to model physical properties of *in vivo* tissues. Existing systems to study confined cell migration largely rely on either ECM-based or rigid materials to provide confinement. The former offer limited ability to isolate variables of matrix stiffness without altering pore size and chemical composition, and the latter fail to mimic situations whereby a cell must generate forces to forge their own path through surrounding tissues (54). Here, we engineer a deformable liquid barrier lending authority to cells themselves in determining the degree of confinement and thus cell morphology. While cell-generated forces control the degree of confinement within a given channel geometry, resistance provided by the environment is tunable by controlling interfacial shape. By allowing such cellular autonomy and tuning environmental confining pressures, interfacial deformations match those of surrounding cells *in vivo*. To date, characterization of leukocyte interaction with surrounding cells *in vivo* is largely absent from existing literature due to limitations in optical access of mammalian models. Here, we validate the physiological relevance of confinement within this system by comparing deformations of the interface to those of surrounding cells during neutrophil interstitial motility within a larval zebrafish model. The ability to capture such interactions *in vivo* introduces an appealing model to study how cellular architecture and other physical properties of surrounding cells themselves regulate the immune response.

Given the temporal changes in tissue architecture that occur throughout aging and many disease states (i.e. tissue wounding, infection, malignancy) (55, 56), understanding how altered tissue mechanics regulates host immune cell motility holds particular relevance to human disease. Within liquid microchannels, pioneer cells that interact with the interfacial barrier require greater interfacial deformations to allow space for migration and as a result migrate slower. Increasing confining pressures by tuning channel geometries negatively impacts motility. These results would predict impaired leukocyte recruitment through tissues composed of stiffer cells, or more tightly packed cellular architectures. Future work might employ genetic engineering tactics to study how altered tissue mechanical properties impact leukocyte motility.

Seminal studies have demonstrated that purely mechanical cues can elicit transitions in cell migration mode. Srivastava et al., and Zatulovskiy et al. found that increasing applied transverse load (*-z* direction) or stiffness of an agarose overlay induced a transition from pseudopodial to bleb-based migration in Dictyostelium cells (12, 57). In such studies, blebs were more often formed at areas of negative membrane curvature (57, 58). In addition, Goudarzi et al., found that blebs formed by zebrafish primordial germ cells occur at regions of local membrane invaginations which act as a source of membrane during bleb expansion (59). In liquid channels, both pioneer and trailing cells are under similar transverse confinement (*z* direction). Yet pioneers interact with the interface providing additional axial resistance (*y* direction). The ability of the interface to respond to cell-generated pressures results in distinct morphological polarity of pioneer cells. It is possible that alterations in membrane curvature induced by pioneer cell interaction with the interface drives transitioning to bleb-based motility. Cells use a number of mechanisms to sense membrane curvature, including BAR-domain proteins and PIEZO channels, both of which have been implicated in regulating bleb formation (57, 59). Our system represents an opportunity to decouple different chemical and mechanical cues (i.e. axial vs. transverse confining pressures) to further investigate the mechanisms by which cells sense their external environment to transition between migration modes.

In addition to applying physiologically-relevant confinement onto migrating cells, the ability to establish and control the shape of a liquid-liquid interface on single-cell length scales represents an intriguing potential to measure spatial and temporal variations in cell generated forces during the course of migration. Where traction force microscopy enables probing forces at the cell-substrate interface, observing the curvature of the liquid interface during cell migration offers a simple readout of transverse forces (exerted in *+z* direction). Distinct from hydrogel-based systems where mechanical properties are complicated by measurement length scale (60), equilibrium pressures of the liquid interface are independent of length scale. Using a read out of interfacial curvature could provide a mathematically straightforward way to measure cell-generated pressures in the transverse direction during motility that is independent of active external intervention required in other methods (i.e. atomic force microscopy).

Thus far, advances in biomaterials engineering have largely been confined to modeling cell interactions with the extracellular matrix. The ability to model physical interactions with surrounding cells provokes an intriguing conceptual shift in the engineering of synthetic materials to model *in vivo* tissues. From both a cell biology and technical engineering perspective, this work advances our understanding of how leukocytes forge paths through dense cellular architectures, while introducing a pioneering technology with implications for further development of soft material-based systems to model an array of *in vivo* tissue properties. Further elucidation of how immune cells sense and respond to physiologically relevant physical cues might facilitate development of therapeutic approaches targeting immune cell migration machinery in the context of human disease.

## Materials and Methods

### Construction of liquid-liquid microchannels

A chambered coverglass (no. 1.5 borosilicate glass, 0.13 to 0.17 mm thick; Thermo Fisher Scientific, 155360) was treated first with O_2_ plasma (Diener Electronic Femto, Plasma Surface Technology) at 60 W for 3 min and then moved to an oven for grafting of PDMS-Silane (1,3-dichlorotetramethylsiloxane; Gelest, SID3372.0; about 10 μl per device) onto the glass surface by chemical vapor deposition (CVD) for 30 mins. The PDMS-grafted slide was masked by a PDMS stamp (stamp construction described in *Li et al*. (47)) and treated with O_2_ plasma at 60 W for 3 min. After surface patterning, the PDMS stamp was removed by tweezers and stored in a clean space for reuse. The chambered coverglass slides were overlaid with oil (silicone oil, 5 cSt; Sigma-Aldrich, 317667) to generate a surface exhibiting ELR to aqueous media. Liquid-liquid channels were created by sweeping a hanging droplet across the patterned surface with a wide-orifice pipette. Channels were constructed with collagen by sweeping a 1:1:1 solution of rat tail collagen I (Corning, 10 mg/mL)) neutralized with one part neutrophil culture media (Roswell Park Memorial Institute (RPMI) 1640 Medium, Thermo Fisher Scientific,11875093) + 2% fetal bovine serum (FBS; Thermo Fisher Scientific, 10437010) and 1% Penicillin/Streptomycin)) and one part 2X HEPES buffer, yielding a final concentration 3.33 mg/mL of rat tail collagen at pH 7.2. For experiments requiring visualization of the collagen layer, one in 15 parts RFP-labeled rat tail collagen was added to allow for confocal imaging of fiber architecture. The collagen solution was incubated at 37 °C, 21% O2, 5% CO2, and 95% RH for 15 minutes to allow for polymerization before transfer to an on-stage incubator (Bold Line, Okolab) for imaging.

### Whole blood collection and neutrophil isolation

All blood samples were drawn according to Institutional Review Boards (IRB) approved protocols per the Declaration of Helsinki at the University of Wisconsin-Madison in the Beebe Lab (IRB# 2020 − 1623) and in the Huttenlocher Lab (IRB#, 2017-003). Informed consent was obtained from all subjects in the study. Whole blood was collected with standard ethylenediaminetetraacetic acid (EDTA) tubes (BD, #36643, EDTA [K2] STERILE, 1054688) and then stored at RT (∼ 22 °C) or 37 °C in stationary storage before isolation.

Neutrophils were isolated from whole blood using magnetic bead-based negative selection per protocol using either the EasySep Direct Human Neutrophil Isolation Kit (STEMCELL, 19666) or the MACSxpress Neutrophil Isolation Kit (Miltenyi Biotech, 130-104-434) purified with the MACSxpress Erythrocyte Depletion Kit (Miltenyi Biotech, 130-098-196). After isolation, the neutrophil pellet was directly resuspended in RPMI 1640 culture media with 2% FBS and 1% Penicillin/Streptomycin (as above). Following resuspension, liquid cultures were stored at 37° in a standard CO_2_ incubator (Thermo Scientific, HERACELL 240i) for no more than 1 hour before seeding into the device and transfer to an onstage incubator for imaging.

### Quantification of cell and channel heights following passive sweep

Primary neutrophils were placed directly into channels by incorporation into the neutralized collagen mixture at concentration of 80k/µL before forming channels by sweep. Cells were labeled immediately after resuspension by incubation at 37 °C and 5% CO_2_ for 15 minutes with with the cytoplasmic dye Calcein-AM green (1 µm; Invitrogen, C3100MP) and the nuclear dye Hoescht 33342 (10µg/mL; Invitrogen, H1399) in RPMI 1640 media with 2% FBS and 1% Penicillin/Streptomycin before combination with the neutralized collagen. The collagen plus cell suspension was swept across the patterned surface as described above to place cells directly into channels. Confocal imaging was performed on a Nikon AR1 spinning disk microscope courtesy of the UW Optical Core to gather z-stacks with step size of 200 nm. Cell and channel heights, in addition to interfacial curvatures, were measured from cross-sectional orthogonal views at regions where a cell was, or was not present (both within the same channel). A single representative cell at the center of the channel was chosen for analysis per unique channel replicate, with a total of three channel replicates for each independent human donor (n = 3).

### Quantification of cell morphology and motility during chemotaxis

To study cell chemotaxis under confinement by the liquid-liquid interface (Fig. 3), channels were constructed by sweeping with neutralized collagen and incubated at 37° to allow polymerization, as described above. Immediately after sweep, excess volume was manually removed by pipette to ensure equal volumes in both inlet and outlet droplets. Following isolation from whole blood, neutrophils were re-suspended in RPMI 1640 media containing 2% FBS and 1% Penicillin/Streptomycin at a concentration of 40,000 cells per µL. In experiments where interfacial height or curvature was measured from volumetric confocal stacks (Fig. 3 and Fig. 4), media was visualized by addition of fluorescein (FITC) sodium salt (40 µM; Sigma Aldrich, F6377). Vybrant DiD (Invitrogen, V22887) was used to label the cell membrane; 5µL dye was added to 1mL of cell suspension in above media immediately following isolation from whole blood (per the Invitrogen protocol) and incubated at 37°C and 5% CO_2_ for 15 minutes prior to washing and cell resuspension to the desired concentration of 40k cells per µL for seeding into the liquid inlet droplets.

Following resuspension, 1.5 µL of cell suspension was added to the inlet droplet by pipette. Immediately after, 1.5 µL of 100 nM *N*-Formylmethionyl-leucyl-phenylalanine (fMLP; Millipore Sigma F3506) in the same media was added to the outlet droplet. After such addition, the cell suspension was mixed by pipetting to ensure homogeneity and cell positioning near the channel entrance. Addition of equal amounts of volume to both the inlet and outlet droplets is essential to restrict generation of convective flow throughout the channel, so that chemokine gradients are driven by diffusion alone. Cell motility was visualized on a Nikon epi-fluorescent microscope (Nikon Eclipse TE3000 with a Nikon Plan Fluor 20x/0.50 objective, motorized stage from Ludl Electronic Products and Prime BSI Express camera from Teledyne Photometrics or a Nikon TI Eclipse inverted microscope with Nikon Plan Fluor 20x/0.50 objective and Hamamatsu camera). Quantification of cell speed was performed in ImageJ/FIJI using the manual tracking plug-in of time lapse movies taken at time intervals of 30 seconds. Volumetric imaging of cells interacting with the interface was performed by labeling the media with FITC dye and imaging with the aforementioned Nikon AR1 spinning disk confocal microscope with step size of 200 nm. Through such volumetric imaging, interfacial heights were measured by the height of the FITC labeled media layer. Similarly, measurements of interfacial pressures during pioneer cell migration (Fig. 4) were made from measuring interfacial curvature (defined by the FITC labeled media) from cross-sectional (*xz* plane) and side (*yz* plane) volumetric images.

### Zebrafish maintenance and handling

Animal care and use was approved by the Institutional Animal Care and Use Committee of University of Wisconsin and strictly followed guidelines set by the federal Health Research Extension Act and the Public Health Service Policy on the Humane Care and Use of Laboratory Animal, administered by the National Institutes of Health Office of Laboratory Animal Welfare. All protocols using zebrafish in this study were approved by the University of Wisconsin-Madison Research Animals Resource Center (protocol M005405-A02). To generate the larval zebrafish model used to visualize neutrophil morphology and keratinocyte deformations, previously published neutrophil-cytoplasm labeled *Tg(LyzC:TagBFP)* fish were crossed with nuclear labeled *Tg(LyzC:H2B-mCherry)* fish to generate a stable line (*Tg(LyzC:TagBFP/LyzC:H2B-mCherry)*. These fish were bred with a stable line expressing cytoplasmic acGFP in basal keratinocytes *Tg(Krtt1c19e:acGFP)*) *(Gift of Dr. Alvaro Sagasti)* to generate larvae used in each experiment. All fish were maintained on the AB background. Following breeding, fertilized embryos were transferred to E3 medium (5 mM NaCl, 0.17 mM KCl, 0.44 mM CaCl_2_, 0.33 mM MgSO_4_, 0.025 mM NaOH, and 0.0003% Methylene Blue) and maintained at 29°C. Larval zebrafish were anesthetized using 0.2 mg/ml tricaine (ethyl 3-aminobenzoate; Sigma-Aldrich) before any experimentation or live imaging.

### Tracking *in vivo* keratinocyte deformations

Dechorionated, tricaine-anesthetized three days-post fertilization (dpf) larvae were mounted in 2% low-melting point agarose (Sigma-Aldrich) on a 35 mm glass bottom dish (#1.5H Glass, Cellvis, CA, USA). Time-lapse imaging of spontaneous neutrophil migration through basal keratinocytes in the yolk sac and neck region was performed on an spinning-disk confocal (CSU-X; Yokogawa) on a Zeiss Observer Z.1 inverted microscope and an electron-multiplying charge-coupled device Evolve 512 camera (Photometrics), with a EC Plan-Neofluar 40×/NA 0.75 air objective (1-2µm optical sections, 2,355 × 512 resolution) using ZenPro 2012 software (Zeiss). Images were captured at 10 second intervals with z-step intervals of 3 µm. Quantification of deformations was performed on resulting maximal intensity projections; background subtraction was manually performed in FIJI/ImageJ in images displayed in the main text (Fig. 5).

### Quantification of bleb formation in primary neutrophils and neutrophil-like PLB-985 cells

Both primary neutrophils and PLB-985 neutrophil-like cells were used to determine the migration mode utilized by neutrophils during interaction with the confining interface (Fig. 6). Immediately following isolation from whole blood, primary neutrophils were incubated with 10 µM CK666 (Millipore Sigma, SML0006) or LY-27632 (Millipore Sigma, Y0503) to inhibit CK-666 or Rho-Kinase at 37 °C and 5% CO_2_ for 30 mins. Bleb-like structures in primary neutrophils were quantified from brightfield time lapse images taken at 1 sec intervals.

PLB-985 cells expressing LifeAct-mRuby were generated by lentiviral transfection. HEK293T cells were grown to 70% confluency in a 10-cm tissue culture dish for each lentiviral target and transfected with pLV-EF1a-IRES-Neo (Addgene #85139) containing LifeAct-mRuby, VSV-G, and CMV 8.9.1. The 72-h viral supernatant was collected and concentrated using a lentivirus concentrator (Lenti-X; Takara Bio Inc.) following the manufacturer’s instructions. PLB-985 cells (1 × 10^6^ total number) were infected with viral supernatant for 3 days in the presence of 15 μg/ml polybrene. Stable cell lines were generated with 1 μg/ml neomycin selection. Resulting PLB-985 cells expressing LifeAct-mRuby were differentiated into neutrophil-like cells by treatment with 1.25% DMSO (Sigma-Aldrich, D2650) for 6 days at 37 °C and 5% CO_2_ in RPMI 1640 with 2% FBS and 1% Penicillin/Streptomycin. Cell motility was assayed in an equivalent assay as that of primary cells, with imaging performed on the previously described Zeiss Observer Z.1 inverted spinning disc confocal microscope, at 0.5-1 sec intervals. Background subtraction was manually performed in FIJI/ImageJ in images displayed in the main text (Fig. 6).

### Statistical Analysis

Data were analyzed (Prism 9.0; GraphPad Software). Statistical significance was assessed using Student’s *t* tests when comparing two conditions/groups, and one-way analysis of variance (ANOVA) corrected using the Tukey’s test when comparing multiple groups.

## Funding

### Biotechnology Training Program

University of Wisconsin-Madison

T23GM135066

PI: Brian Fox

### Ruth L. Kirschstein National Research Service Award (NRSA) Fellowship

National Heart, Lung, and Blood Institute

1F30HL174128

PI: Jon Schrope

### Carbone Cancer Center Support Grant

University of Wisconsin NIH P30CA014520

### Microscale models of inflammation and its resolution

NIH R01AI34749

PI: Beebe and Huttenlocher

### Cell Migration and Wound Repair

NIH R35GM118027-08

PI: Huttenlocher

### A multiplexed micro scale assay for real time analysis of pediatric immune cell function

NIH U24AI152177

PI: Beebe

### Under-oil open microfluidic system (UOMS) for studying systemic fungal infection

NIH R01AI54940

PI: Beebe

## Data and materials availability

All data are available in the main text or the supplementary materials, or upon request.

## Acknowledgments

The authors would like to thank the University of Wisconsin Imaging Core for the use of microscopy facilities as well as Biorender and Imaris for use of software to make figure schematics.

