## Supplementary Figures for "Confinement by liquid-liquid interface replicates *in vivo* neutrophil deformations and elicits bleb based migration"

### Supplemental Information for

#### Liquid-liquid interfaces enable tunable cell confinement to recapitulate cell deformations during neutrophil interstitial migration *in vivo*

Jonathan H. Schrope[1,2,3,4], Adam Horn[2], Kaitlyn Lazorchak[2,4] , Clyde W. Tinnen[3], Jack J Stevens[1,2], Mehtab Farooqui[3,5], Tanner Robertson[2], Jiayi Li[1], David Bennin[5], Terry Juang[1,3], Adeel Ahmed[5], Chao Li[5,7], Anna Huttenlocher[2,6,7,*], David J Beebe[1,3,5,7,*]

[1]Department of Biomedical Engineering, University of Wisconsin-Madison, Madison, WI, USA.
[2]Department of Medical Microbiology and Immunology, University of Wisconsin-Madison, Madison, WI, USA.
[3]Department of Pathology and Laboratory Medicine, University of Wisconsin-Madison, Madison, WI, USA.
[4]Medical Scientist Training Program, University of Wisconsin-Madison, Madison, WI, USA.
[5]Carbone Cancer Center, University of Wisconsin-Madison, Madison, WI, USA.
[6]Department of Pediatrics, University of Wisconsin-Madison, Madison, WI, USA.
[7]These authors contributed equally.

*** Co-corresponding authors**: Anna Huttenlocher, David J Beebe

**This PDF file includes:**

Supporting Text
Figures S1 to S9
Legends for Movies S1 to S7
SI References

**Other supporting materials for this manuscript include the following:**

Movies S1 to S7

#### Supporting Information Text

##### Results

##### The liquid-liquid interface is sufficiently rigid to control cell positioning within a chemical gradient

As neutrophils mechanically interact with surrounding cells in the body, chemokine gradients arising from sites of injury or infection regulate motility to sites of inflammation. We next sought to generate a chemotaxis assay under cell confinement by a liquid interface, which would require spatially patterning of cells within a chemokine gradient. Such gradient generation and control of cell positioning (i.e. "cell trapping") is commonly achieved using rigid materials that neither cells nor chemokine can permeate (1–3). Given that the liquid-liquid interface exerts pressures sufficient to confine cells, we hypothesized that this interface could act as a physical barrier to spatially trap cells at the channel entrance within a chemical gradient diffusing from the outlet droplet. Thus, channels were constructed with collagen, followed by sequential addition of cells to the inlet droplet by direct pipetting (**Fig. S6A**). The result is a layer of media upon a layer of fibrillar collagen (**Fig. S6B**). If the oil-media interface possesses sufficient rigidity to spatially trap cells within the media layer, then the media layer height would control cell positioning along the channel entrance and thereby enable control of positioning within the chemokine gradient. We find that cells become spatially trapped at the entrance to the channel where interfacial height is comparable to neutrophil diameter (~ 6-12 μm (4)) (**Fig. S6C**). Addition of FITC dye to the outlet droplet establishes a chemical gradient that is stable over experimental timescales (>3 hours) (**Fig. S6D-E**). Taken together, these results demonstrate a novel method of cell trapping within open microscale systems using a liquid-liquid interfacial barrier to control cell positioning within a chemical gradient to study chemotaxis under confinement.

#### Supplementary Figures

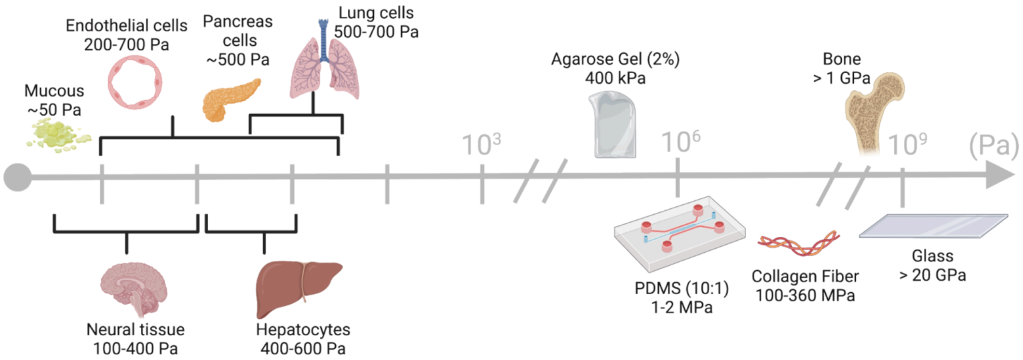


**Figure S1. Review of elastic moduli of individual cells.** Commonly used *in vitro* materials exhibit elastic moduli orders of magnitude higher than that of single cells. Stiffness values depicted here gathered from a review by Guimarães et al that compiled studies reporting elastic moduli of single cells (5).

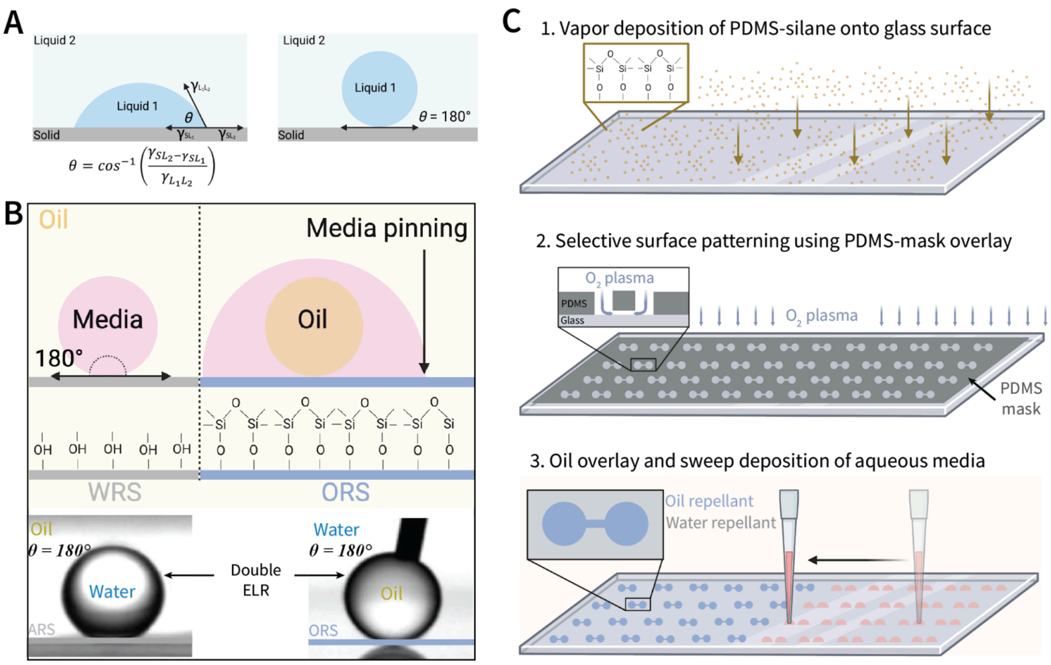


**Figure S2. Construction of liquid microchannels by selective patterning of a glass surface to exhibit double ELR. A)** The degree to which a solid surface repels a liquid within a solid-liquid-liquid three-phase system is reflected in the Young's contact angle ($\theta$) that a liquid droplet (Liquid 1) makes on the surface. The Young's contact angle is determined by the balance between all interfacial energies ($\gamma_{SL1}$, $\gamma_{SL2}$, $\gamma_{L1L2}$) present within the system. Precise engineering of this energy balance can result in a surface exhibiting Exclusive Liquid Repellency (ELR), characterized by a Young's contact angle of 180°. **B)** Selective chemical patterning of a glass surface to exhibit double ELR results in regions ELR to aqueous media in the presence of oil (Aqueous Repellent Surface, ARS) or ELR to oil in the presence of media (Oil Repellent Surface, ORS). **C)** Differential surface pattering is obtained by grafting liquid PDMS-silane onto a virgin glass surface by vapor deposition to generate a homogenous ARS surface (repellent to aqueous media in the presence of an oil overlay). A PDMS-based mask is placed on top of the slide and treated with O2 plasma. Surface treatment is etched away in areas exposed to $O_2$ plasma, where non-exposed areas remain ELR to aqueous media. The mask is removed, and the surface overlaid with oil to generate a double ELR surface, that is, regions repellent to aqueous media (ARS) and regions repellent to oil (ORS).

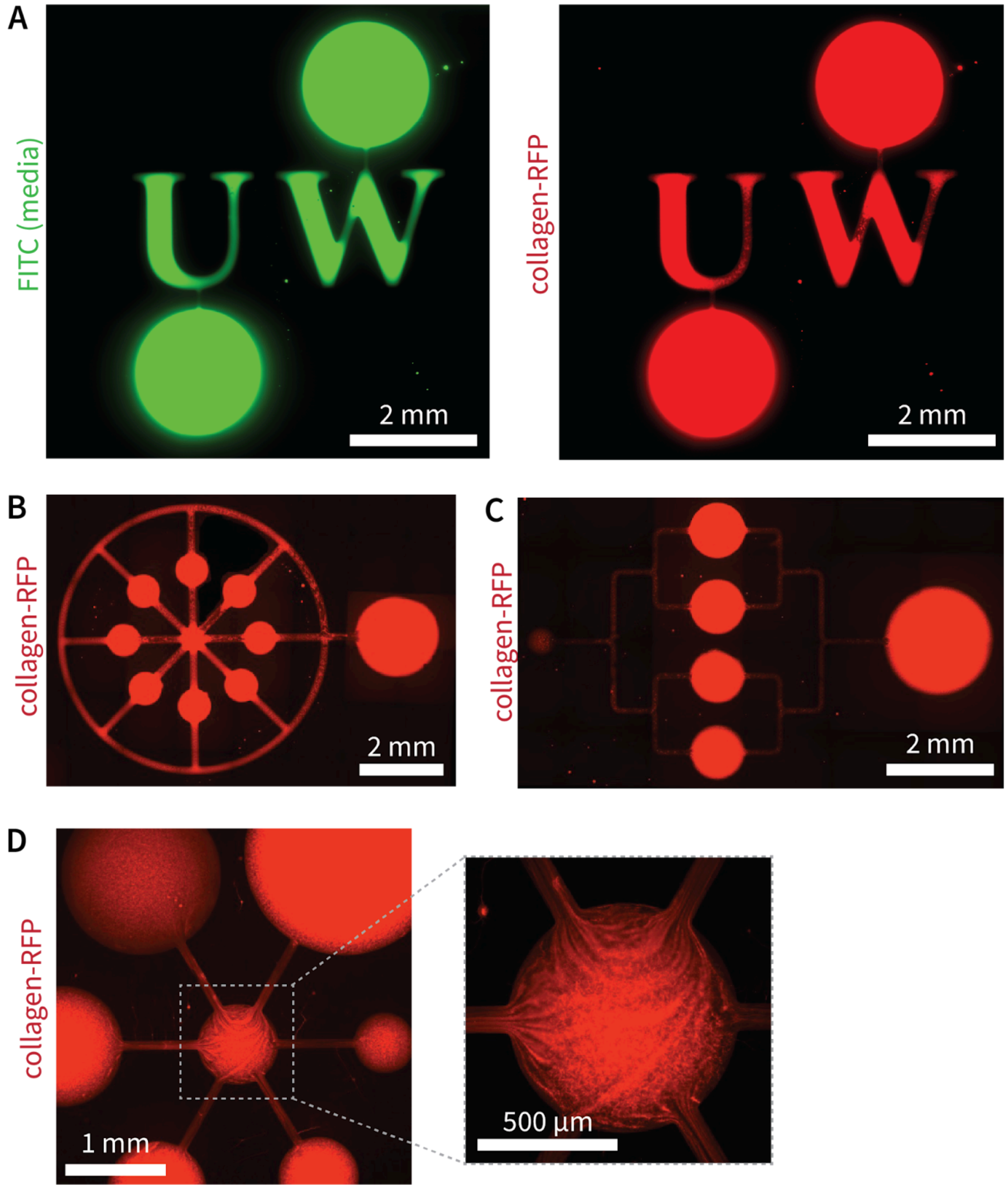


**Figure S3. Examples of different patterns of liquid channels. A)** Representative images of the letters “UW.” Left shows an image of aqueous media labeled with FITC dye and right shows the same pattern made of RFP-labeled fibrous collagen. **B-D**) Different patterns of collagen channels.

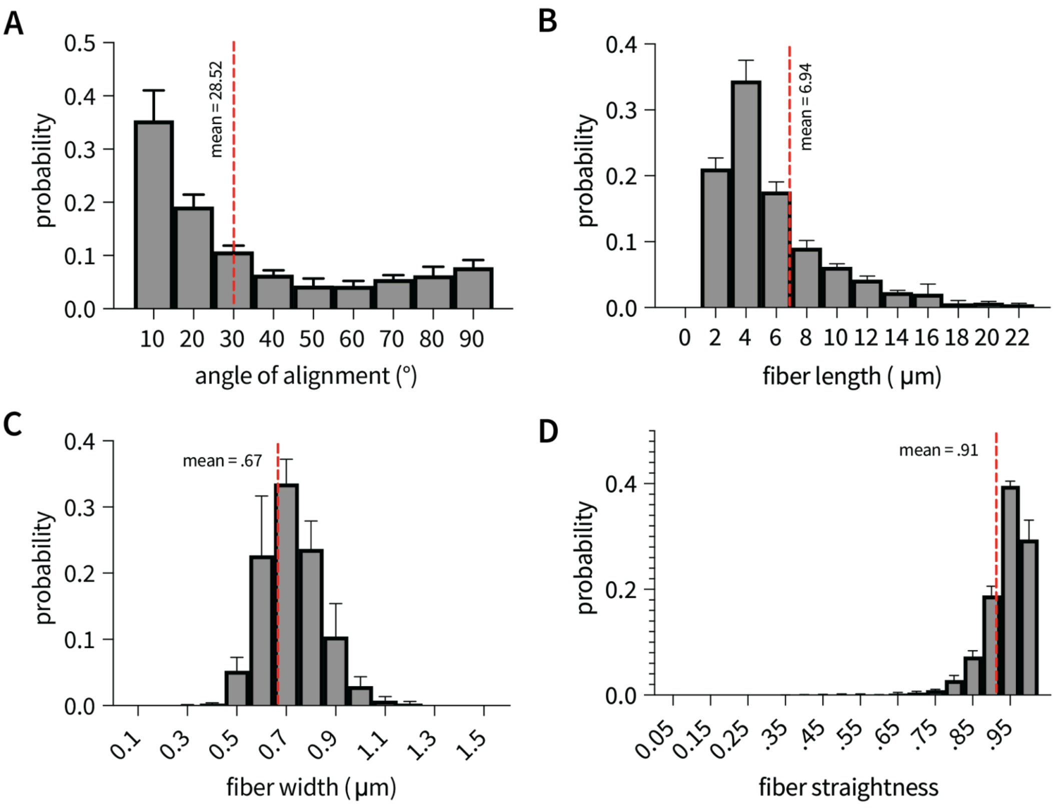


**Figure S4. Characteristics of collagen coating.** The mean **A)** angle of alignment **B)** fiber length **C)** fiber width and **D)** fiber straightness were calculated over 8 channels (30 µm width) analyzed using CurveAlign software (developed by LOCI at UW-Madison). Notably, collagen structure was not altered during nor after neutrophil migration (data not shown).

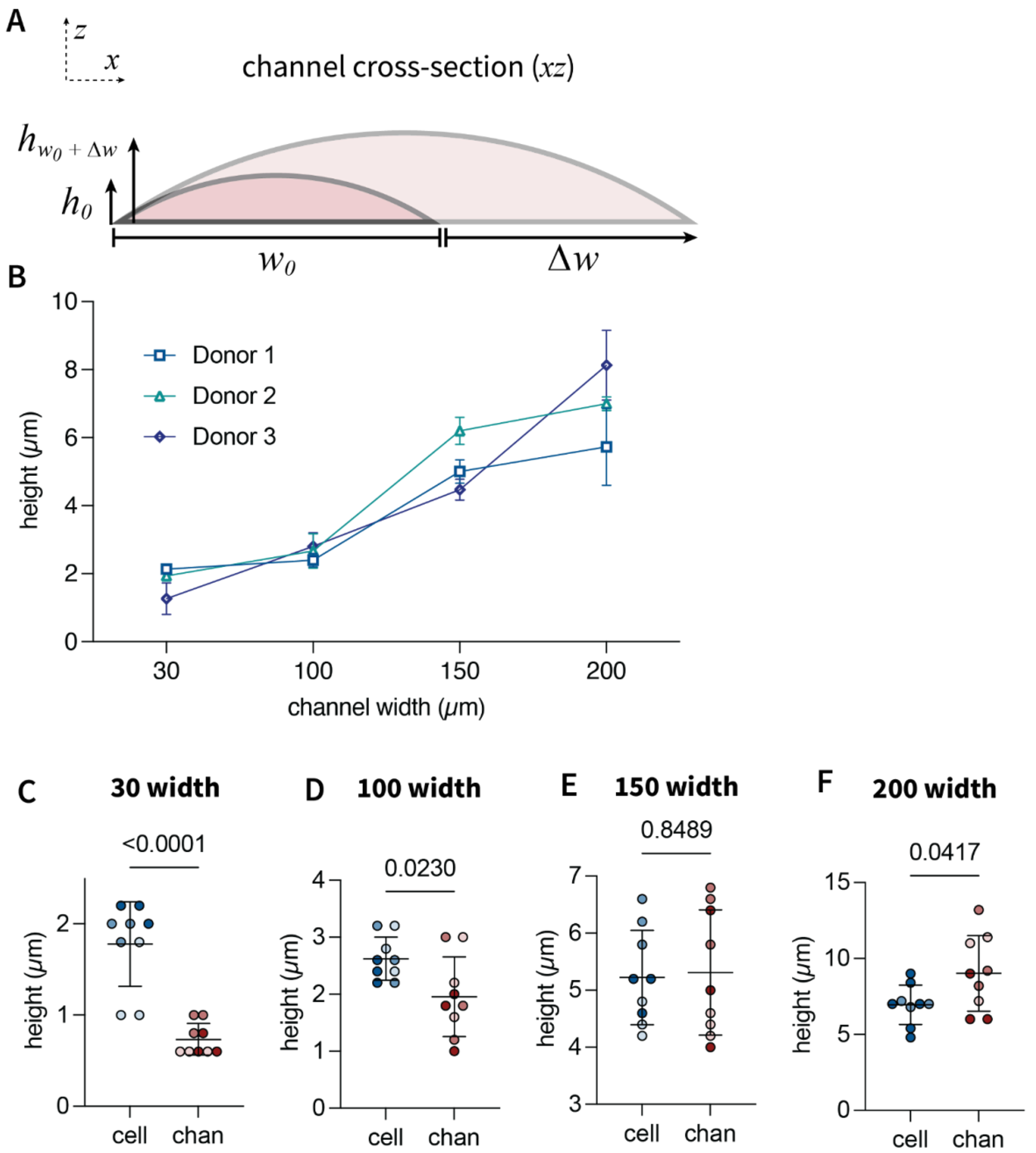


**Figure S5. Channel and cell heights following direct placement of immoble cells into channels. A)** Schematic depicting how increasing channel width increases height, measured as the maximum height at the center of the channel. **B)** Plot of cell heights including donor variability following direct incorporation into channels of varying width by sweep technique. Each point represents the mean height of a cell (n = 3) measured on independent channel replicates (n = 3) for each donor. **C-F)** Plots comparing cell height to channel height (in areas absent of cells) on channels of varying width. On small channels with heights significantly less than that of cells (30 and 100 μm width), cells deform the interface to generate height greater than initial channel heights. On large channels (i.e. 200 μm width), cells assume a natural (i.e. non-confined) diameter less than that of the channel height. Each point represents a single cell, shade represents independent donor (n = 3). Statistical significance was determined by an unpaired t-test assuming equal variance.

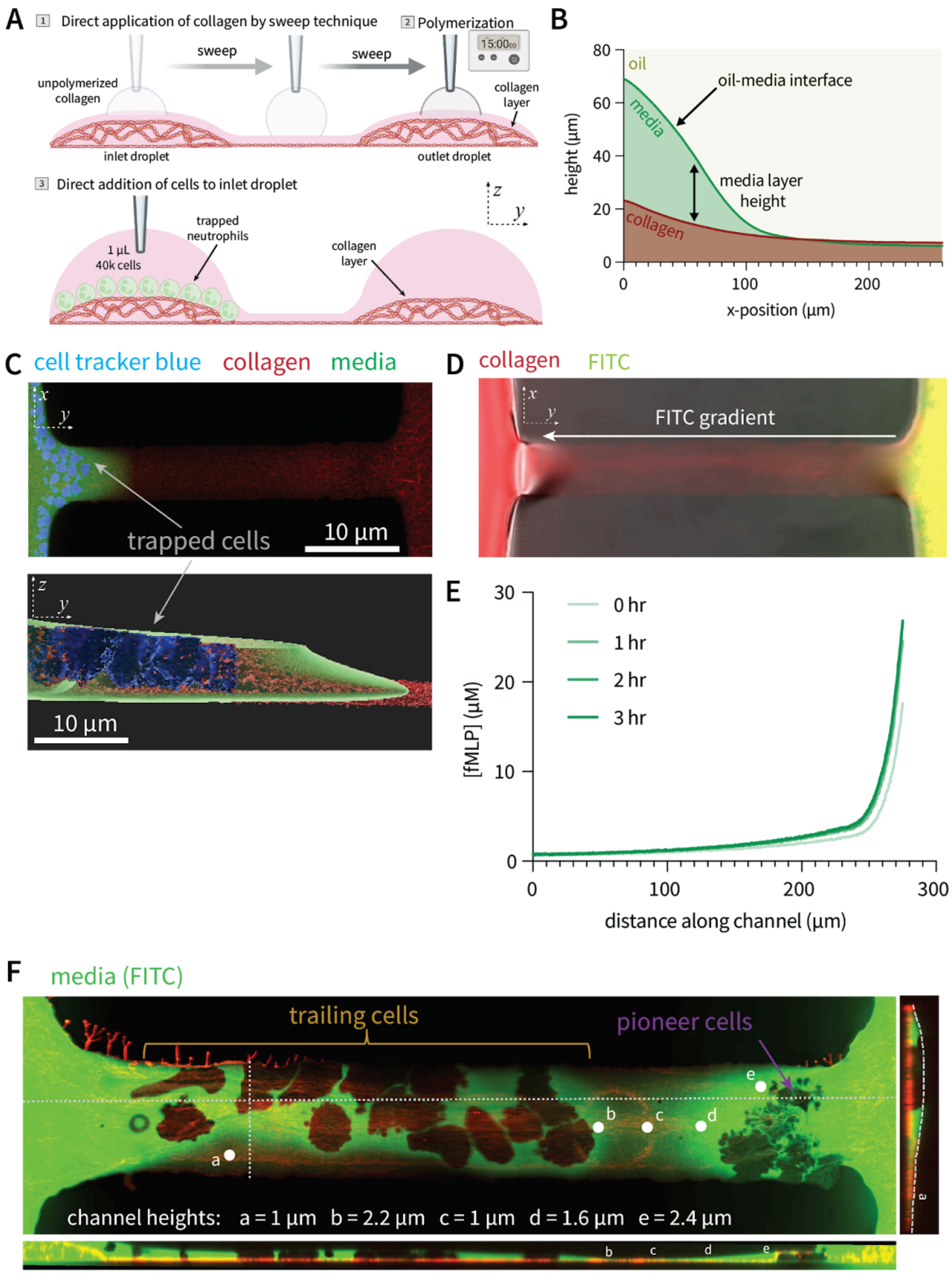
A
1 Direct application of collagen by sweep technique
2 Polymerization
15:00
sweep
sweep
unpolymerized collagen
collagen layer
inlet droplet
outlet droplet
3 Direct addition of cells to inlet droplet
trapped neutrophils
1 µL 40k cells
collagen layer
z
y
B
oil
oil-media interface
media
media layer height
collagen
height (µm)
x-position (µm)
C
cell tracker blue collagen media
x
y
trapped cells
10 µm
z
y
10 µm
D
collagen FITC
x
y
FITC gradient
E
0 hr
1 hr
2 hr
3 hr
[fMLP] (µM)
distance along channel (µm)
F
media (FITC)
trailing cells
pioneer cells
a
b
c
d
e
channel heights: a = 1 µm b = 2.2 µm c = 1 µm d = 1.6 µm e = 2.4 µm

**Figure S6. Single cell trapping and gradient formation within liquid channels. A)** Schematic depicting cell trapping assay whereby cells are added to the inlet droplet following creation of collagen-coated channels. **B)** Representative profile of the media and collagen layers for a 200 µm width channel. The height of the media layer converges to zero at some point along the length of the channel (~ 130 µm for a 200 µm width channel). **C)** Representative confocal top and side (Imaris reconstruction) view of cells at the entrance of a 30 µm width and 300 µm length channel spatially trapped at the entrance where media layer height is comparable to cell height (~ 8 µm for neutrophils). **D-E)** Addition of FITC to the outlet droplet establishes a chemical gradient that is relatively stable over experimental timescales of ~ 3 hours. Approximate concentration of fMLP along channel length was extrapolated from measurements of FITC gradients over time (similar molecular weights of FITC 389.382 g/mol vs fMLP 437.56 g/mol) **F)** Representative confocal image of migratory primary neutrophils under confinement by the liquid-liquid interface. Media is visualized by FITC dye added in equal concentrations (40 µM) to the inlet and outlet droplets.

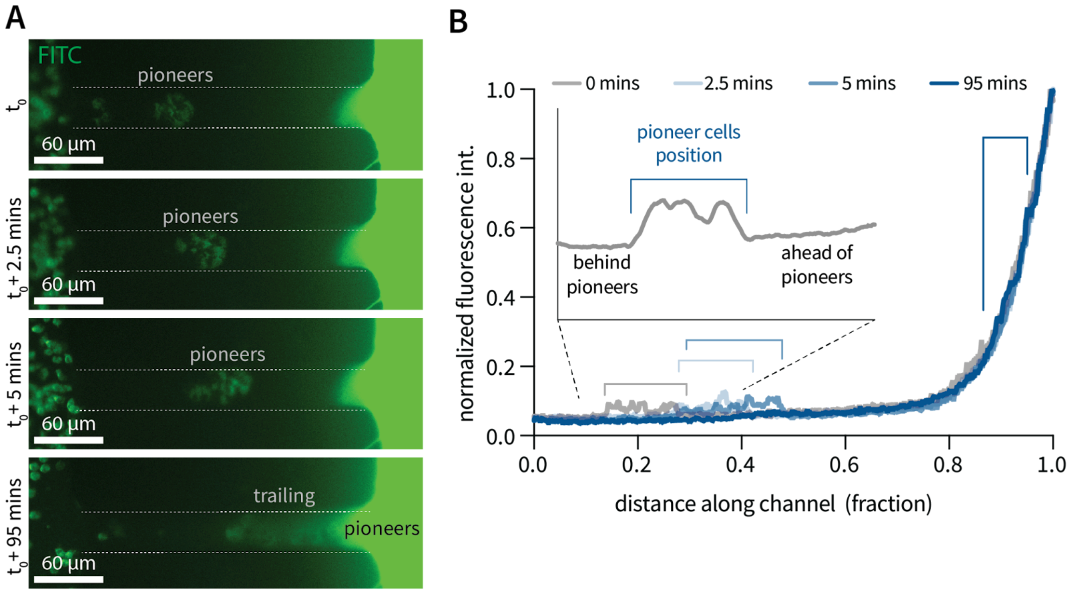


**Figure S7. Gradient stability during neutrophil migration. A)** Representative timelapse images of gradient characterization during cell migration using FITC dye added to the outlet droplet (MW = 389.382 g/mol) as a proxy for fMLP chemoattractant (MW = 437.56 g/mol). **B)** Under this method, cells are partially labeled by FITC to mark the position of pioneer cells within the device. The FITC gradient in front of, or behind the cells is unperturbed over time.

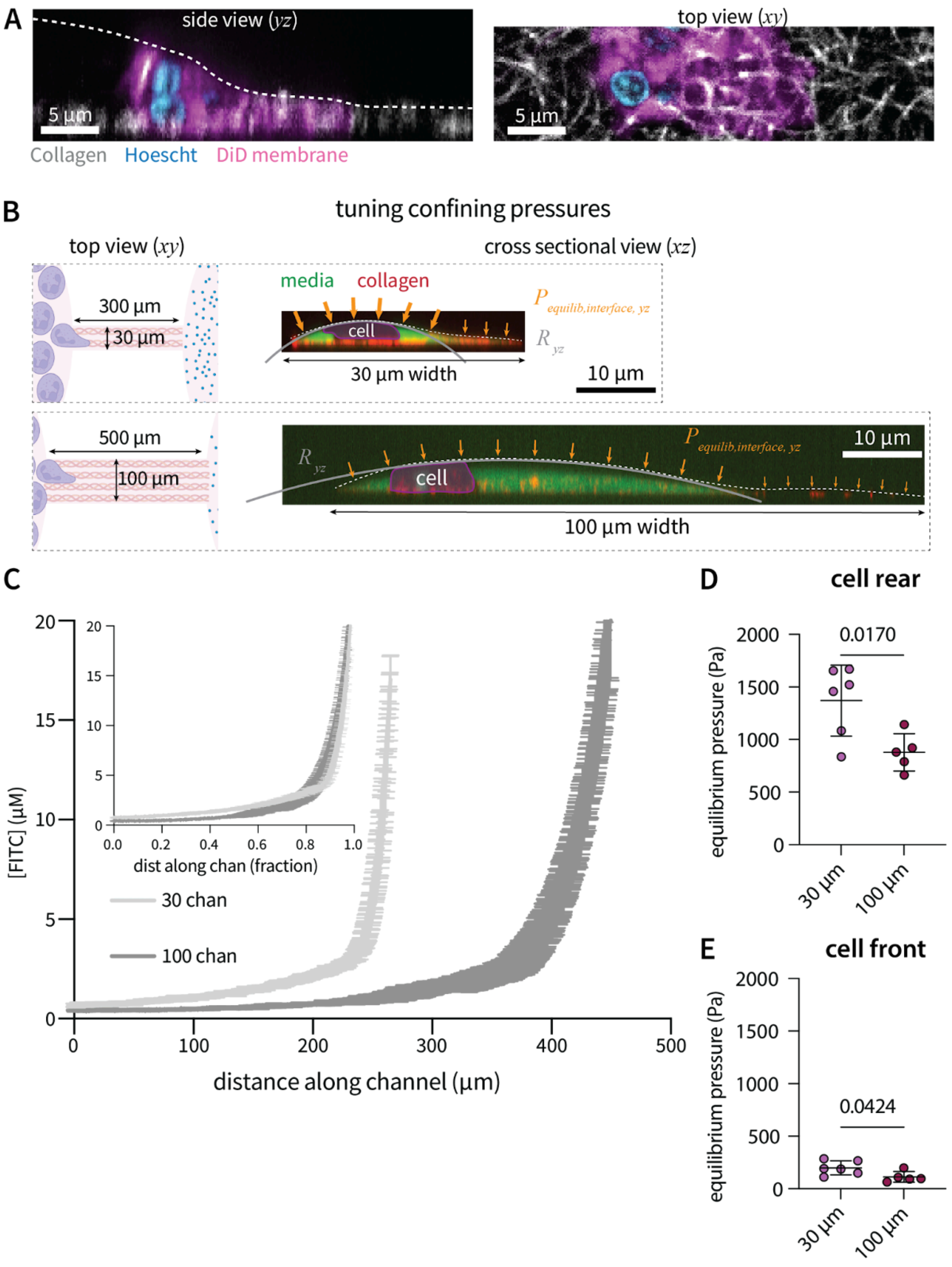
A
side view (yz)
5 µm
top view (xy)
5 µm
Collagen Hoescht DiD membrane
B
tuning confining pressures
top view (xy)
cross sectional view (xz)
300 µm
30 µm
media collagen
cell
30 µm width
P equilib,interface, yz
R yz
10 µm
500 µm
100 µm
cell
100 µm width
10 µm
C
[FITC] (µM)
distance along channel (µm)
dist along chan (fraction)
30 chan
100 chan
D
cell rear
equilibrium pressure (Pa)
0.0170
30 µm
100 µm
E
cell front
equilibrium pressure (Pa)
0.0424
30 µm
100 µm

**Figure S8. Characterization of cell morphology as a function of channel width A)** Confocal side and top view of representative pioneer cell on a 100 μm channel width on a collagen substrate (gray). **B)** Schematic and cross-sectional views of cell height on channels of differential geometry. **C)** Tuning channel geometries does not significantly alter the chemical gradient (all migration data analyzed up to .8 fraction along length of channel). The 30 μm width 300 μm length channel yields a slightly steeper gradient which would hypothetically yield faster migration (our data indicates slower, attributed to differences in confining pressures). **D-E)** Quantification of equilibrium confining pressures over the rear (D) and front (E) of migratory cells within channels of different width. Statistical significance determined by an unpaired, two-sample t-test assuming equal variance.

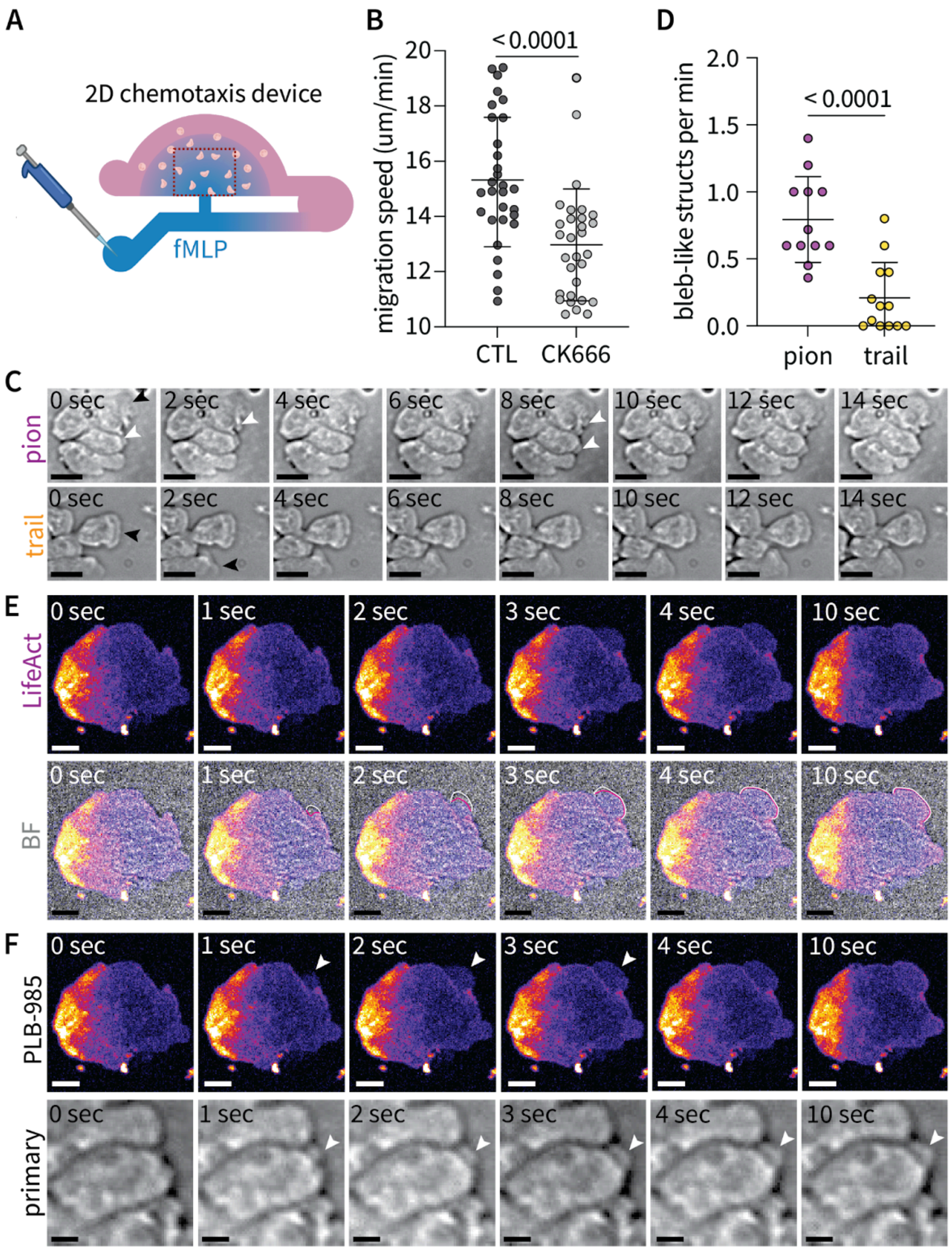
A
2D chemotaxis device
fMLP
B
< 0.0001
migration speed (um/min)
20
18
16
14
12
10
CTL
CK666
D
< 0.0001
bleb-like structs per min
2.0
1.5
1.0
0.5
0.0
pion
trail
C
pion
trail
0 sec
2 sec
4 sec
6 sec
8 sec
10 sec
12 sec
14 sec
E
LifeAct
BF
0 sec
1 sec
2 sec
3 sec
4 sec
10 sec
F
PLB-985
primary

139

**Figure S9. Characterization of primary neutrophil and neutrophil-like PLB-985 cell migration mode. A)** Schematic of two-dimensional chemotaxis device **B)** Pioneer cell migration speed in the absence of confinement is decreased under treatment with Arp-2/3 inhibitor CK666. Significance was determined by an independent two sample t-test between the two experimental groups (n = 30 cells each, 10 tracked over 3 device replicates) **C)** Brightfield time lapse images of pioneer and trailing cell migration. White arrows indicate bleb-like structures and black arrows indicate sheet-like pseudopodial protrusions. Scale bar represents 10 μm. **D)** Quantification of bleb-like structures per minute for pioneer and trailing primary neutrophils. Circles represent individual cells, data pooled evenly over three independent donor replicates with three independent channels per donor replicate. **E)** Timelapse images of LifeAct-mRuby expressing neutrophil-like PLB-985 cells depicting membrane extension (white line) past actin signal (red line) at initial stages of bleb formation. Scale bars represent 5 μm. **F)** Timelapse images depicting similar shape and formation kinetics between bleb-like structures of primary neutrophils and blebs of neutrophil-like PLB-985 cells. Scale bars represent 5 μm.

#### Supplementary Movie Legends

**Movie S1 (separate file). Liquid Channels.** Three-dimensional artistic rendering of collagen-coated liquid channels located between two media droplets (inlet and outlet). Mesh denotes oil-media interface.

**Movie S2 (separate file). Pioneer cell morphology.** Three-dimensional confocal reconstruction of a pioneer cell during migration, generated using Icy image analysis software.

**Movie S3 (separate file). *In vivo* neutrophil interstitial migration.** Timelapse of neutrophil interstitial migration depicting deformations of surrounding basal keratinocyte cells (green) during the protrusion stage and passage of the cell body (blue) containing the nucleus (red). Time given as mm:ss.

**Movie S4 (separate file). Pioneer and trailing primary neutrophil migration.** Timelapse depicting migration of pioneer (left) and trailing (right) primary neutrophils. White arrows depict rapid bleb-like protrusions and black arrows sheet-like pseudopodial protrusions. Images taken at an interval of 0.5 sec, time given as ss:ss.

**Movie S5 (separate file). Trailing PLB-985 cell LifeAct timelapse.** Timelapse depicting sheet-like protrusions of a trailing PLB-985 cell expressing LifeAct-mRuby migrating within a 30 μm width channel.

**Movie S6 (separate file). Pioneer PLB-985 cell LifeAct timelapse.** Timelapse depicting bleb protrusions of a pioneer PLB-985 cell expressing LifeAct-mRuby migrating within a 30 μm width channel.

**Movie S7 (separate file). Transition to blebbing.** Timelapse depicting cell transition from sheet-like pseudopodia to blebs upon reaching the interface in LifeAct-mRuby expressing PLB-985 cells within 30 μm width liquid channels. Movie contains a pioneer cell (top) and trailing cell (bottom) that migrates fast enough to reach the interface and transition to bleb protrusions at the leading edge. Images taken at 1 sec intervals.

#### Supplementary Information References
